# eIF4A1 is essential for reprogramming the translational landscape of Wnt-driven colorectal cancers

**DOI:** 10.1101/2023.11.10.566546

**Authors:** Joseph A. Waldron, Georgios Kanellos, Rachael C. L. Smith, John R. P. Knight, June Munro, Constantinos Alexandrou, Nikola Vlahov, Luis Pardo-Fernandez, Madeleine Moore, Sarah L. Gillen, Douglas Strathdee, David Stevenson, Fiona C. Warrander, Kathryn Gilroy, Colin Nixon, Barbara Cadden, Ian Powley, Leah Officer-Jones, Fiona Ballantyne, Jennifer Hay, Kathryn Pennel, Joanne Edwards, Andrew D. Campbell, Rachel A. Ridgway, Seth B. Coffelt, Jim Norman, John Le Quesne, Martin Bushell, Owen J. Sansom

## Abstract

Dysregulated translation is a hallmark of cancer. Targeting the translational machinery represents a therapeutic avenue which is being actively explored. eIF4A inhibitors target both eIF4A1, which promotes translation as part of the eIF4F complex, and eIF4A2, which can repress translation via the CCR4–NOT complex. While high eIF4A1 expression is associated with poor patient outcome, the role of eIF4A2 in cancer remains unclear. Furthermore, the on-target toxicity of targeting specific eIF4A paralogues in healthy tissue is under-explored. We show that while loss of either paralogue is tolerated in the wild-type intestine, eIF4A1 is specifically required to support the translational demands of oncogenic Wnt signalling. Intestinal tumourigenesis is suppressed in colorectal cancer models following loss of eIF4A1 but accelerated following loss of eIF4A2, while eIF4A inhibition with eFT226 mimics loss of eIF4A1 in these models.

## Main

Mutation of the *APC* tumour suppressor gene is the most common genetic driver alteration in colorectal cancer (CRC), leading to ligand-independent Wnt signalling and activation of TCF/LEF transcriptional targets, e.g., *AXIN2*, *MYC,* and *CCND1* (*1*). Many of the key processes driven by *Apc* mutation *in vivo*, e.g., hyperproliferation and perturbed differentiation, have been shown to be dependent on Myc (*2*). However, the mechanisms by which the transcriptome is co-ordinately converted to the proteome are not well defined. The RNA regulon model (*3–5*) has been proposed to describe how post-transcriptional control of functionally related mRNAs are coordinated, offering a target for therapeutic intervention in cancer. Indeed, oncogenic signalling in CRC critically reprogrammes translational control (*6*), and targeting translation is actively being explored as a potential therapeutic avenue in CRC and other cancer settings (*7–10*). Translation initiation is regarded as the rate-limiting step for most mRNAs and, importantly, the recruitment and positioning of the initiating ribosome on mRNAs is dependent on components of the mRNA cap-binding complex, eukaryotic initiation factor (eIF) 4F (*11, 12*), which lies at the nexus of multiple mitogenic signalling pathways (*13*). The catalytic component of this complex, eIF4A1, is a DEAD-box RNA helicase, thought to be predominantly required for unwinding secondary structures within 5’UTRs (*14–16*), while also possessing a helicase-independent role in ribosome recruitment (*17, 18*). However, two highly similar paralogues of eIF4A, eIF4A1 and eIF4A2, are involved in translation (*19, 20*). Until recently these two paralogues were considered redundant, as they share roughly 90% amino acid identity and behave almost identically *in vitro* (*19*). Despite these similarities, they have distinct tissue expression patterns (*21, 22*) and while eIF4A2 is able to promote the translation of specific mRNAs as part of the eIF4F complex in cells with higher expression of eIF4A2 relative to eIF4A1 (*23, 24*), it can also function as part of the CCR4-NOT complex to repress translation (*23, 25, 26*). Furthermore, eIF4A1 expression has been shown to associate with poor survival in several cancers (*27–32*), whereas the association of eIF4A2 expression with survival is much less clear and varies between different cancers (*33–35*). Several compounds targeting eIF4A have shown promising results in mouse models (*36–42*) and, more recently, in clinical trials (*43, 44*), but these target both paralogues (*45–47*). It is therefore critical that we better understand the effects and implications of eIF4A inhibition on tumourigenesis.

### Targeting *Eif4a1* or *Eif4a2* does not disrupt intestinal homeostasis

To better understand the functional roles of eIF4A1 and eIF4A2 in intestinal homeostasis, we examined their expression in wild-type (WT) mouse intestinal tissue and revealed that eIF4A1 and eIF4A2 exhibit distinct expression patterns (Fig. 1A). Expression of eIF4A1 is localized mainly to intestinal crypts, the highly proliferative, stem cell–containing compartment of the intestinal epithelium, extending partially upwards along the length of the villi (Fig. 1A-B). However, eIF4A2 expression is predominantly in differentiated villus cells, with its expression in crypts restricted to the secretory Paneth cells (Fig. 1A). This is consistent with previous observations that eIF4A1 and eIF4A2 are expressed mainly in proliferative and differentiated/secretory cells, respectively (*21, 22*). Given their distinct expression patterns, we expect each paralogue to be essential for the translational landscape of their respective expressing intestinal cells.

**Fig. 1.**
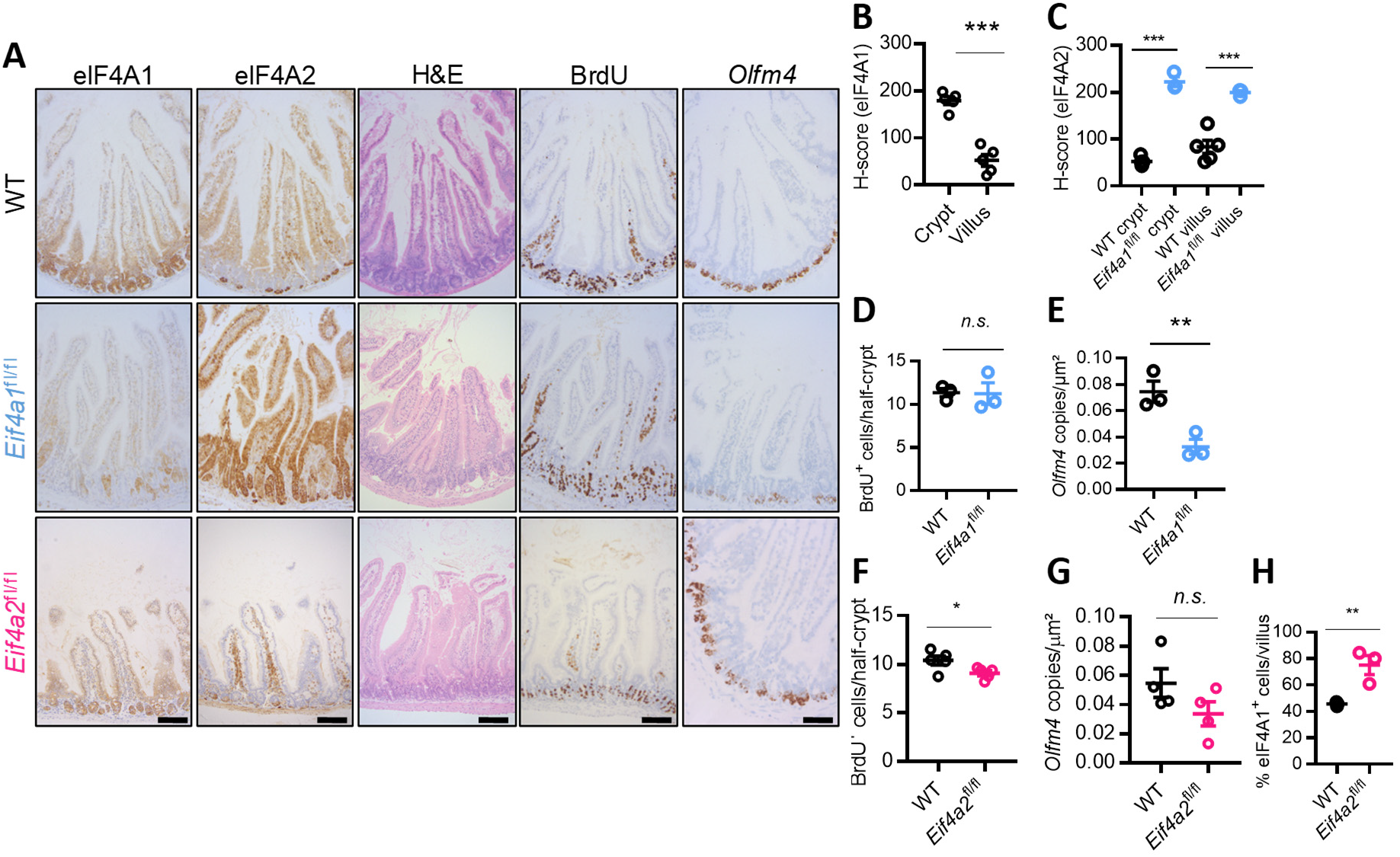
eIF4A1 and eIF4A2 are differentially expressed within the mouse small intestine and can be targeted without disrupting intestinal homeostasis. **(A)** Representative staining for eIF4A1, eIF4A2, H&E, BrdU, and *in situ* hybridisation (ISH) for *Olfm4* in sections of small-intestinal epithelium from *VillinCreER^T2^* (WT, n=5), *VillinCreER^T2^ Eif4a1*^fl/fl^ (*Eif4a1*^fl/fl^, n=5), and *VillinCreER^T2^ Eif4a2*^fl/fl^ (*Eif4a2*^fl/fl^, n=5) mice four days post tamoxifen injection. Scale bars, 100 μm. **(B)** HALO H-score quantification of eIF4A1 expression in the crypts and villi of the murine WT small intestine. n=5 WT mice; p<0.0001. **(C)** HALO H-score quantification of eIF4A2 expression in crypts and villi from WT and *Eif4a1*^fl/fl^ small intestines from (A). WT (crypt), n=5 vs *Eif4a1*^fl/fl^ (crypt), n=3; p<0.0001. WT (villus), n=5 vs *Eif4a1*^fl/fl^ (villus), n=3; p<0.0001. **(D)** Quantification of BrdU⁺ cells in WT and *Eif4a1*^fl/fl^ small intestines from (A). n=3 per genotype; p=0.35. **(E)** HALO quantification of *Olfm4* ISH staining in *Eif4a1*^fl/fl^ and WT small intestines. n=3 per genotype; p=0.0066. **(F)** Quantification of BrdU⁺ cells WT and *Eif4a2*^fl/fl^ small intestines from (A). n=5 per genotype; p=0.039. **(G)** HALO quantification of *Olfm4* ISH staining in WT and *Eif4a2*^fl/fl^ small intestines. n=4 per genotype; p=0.15. (H) HALO quantification of eIF4A1 positively stained cells per villus in WT vs *Eif4a2*^fl/fl^ small intestine. n=3 per genotype; p=0.0077. All data are presented as mean ± SEM and were statistically assessed by unpaired two-tailed t-tests, except in (C) where one-way ANOVA and Tukey’s multiple comparisons test were used. *p <0.05, **p <0.01, *** p <0.001. n.s., not significant.

Unexpectedly, conditional homozygous loss of either paralogue in the intestine, was well tolerated in the short term (Fig. 1A and fig. S1A-D). Loss of eIF4A1 was accompanied by a striking upregulation in eIF4A2 expression, including a spatial expansion into the crypts (Fig. 1C). Loss of eIF4A1 did not overtly impact intestinal architecture or proliferative capacity (Fig. 1A, D and fig. S1B-C), although we did observe an acute reduction in stem cell numbers (Fig. 1E), while no changes were observed in the Paneth or goblet cell populations (fig. S1A). However, *Eif4a1*^fl/fl^ mice had recovered eIF4A1 expression along their intestine by 30 days post induction, together with stem cell populations, indicating that escaper cell populations that retain eIF4A1 expression have a competitive advantage against eIF4A1-null cells (fig. S1E). This suggests that eIF4A2 can only partially compensate for the loss of eIF4A1.

*Eif4a2* could be targeted with no disruption to intestinal homeostasis (Fig. 1A, F-G), and loss of eIF4A2 was also accompanied by an expansion of the eIF4A1 expression domain to the intestinal villi (Fig. 1H). Differentiated and secretory Paneth cells, which predominantly express eIF4A2, were unaffected (fig. S1A). Ablation of eIF4A2 expression was maintained long term (fig. S1F), suggesting that loss of eIF4A2 can be tolerated, presumably through the compensatory expression of eIF4A1.

Notably, concurrent deletion of both *Eif4a* paralogues led to gut toxicity with disruption of the tissue architecture, abrogation of cell proliferation, and complete loss of intestinal stem cells (fig. S1G). Overall, these data suggest functional redundancy between eIF4A1 and eIF4A2 in the intestine of WT mice, with individual gene deletion eliciting compensatory expression of the opposing paralogue. However, whereas loss of eIF4A2 is maintained long-term, eIF4A1-positive cells out-compete eIF4A1-null cells. This suggests that intestinal cells, particularly the stem cells, are more reliant on eIF4A1 to support their translational landscape. In support of this, *Eif4a1*^fl/fl^ *Eif4a2^fl/+^* mice, in which the elevated ectopic eIF4A2 expression induced following *Eif4a1* loss is dampened, display gut toxicity, whereas, a single copy of *Eif4a1* alongside homozygous loss of *Eif4a2* is enough to maintain gut homeostasis long-term (fig. S1G).

### eIF4A1, but not eIF4A2, is required for intestinal proliferation following acute loss of *Apc*

The finding that eIF4A1 loss selectively depleted the Wnt-dependent stem cell populations, without impacting short-term gut homeostasis, suggested that eIF4A1 is essential for their maintenance. Hence, we sought to determine the effect of the loss of either eIF4A1 or eIF4A2 on the hyperproliferative crypt progenitor phenotype, driven by acute homozygous *Apc* loss and consequent hyperactivation of the Wnt-signalling pathway, within intestinal crypts (*48*). Loss of eIF4A1 in *VillinCre^ER-T2^ Apc*^fl/fl^ (hereafter *Apc*^fl/fl^) mice was again accompanied by an expansion of the eIF4A2 expression domain into the crypts (Fig. 2A). However, despite the compensatory increase in eIF4A2, loss of eIF4A1 significantly suppressed crypt hyperproliferation in both the small intestine and the colon (Fig. 2A-B and fig. S2A-B). This reduction in proliferation in *Apc*^fl/fl^ crypts to WT levels (compare Fig. 1D with 2B), following the loss of eIF4A1, is a phenocopy of the loss of Myc (*2*). This suggests that eIF4A1 is required to reprogram the translational landscape following *Apc* loss, akin to the dependency of the transcriptional landscape on Myc. Loss of eIF4A2 led to only a modest reduction in proliferation in the small intestine of *Apc*^fl/fl^ mice, which was significantly smaller than that elicited through loss of eIF4A1 (Fig. 2A-B) and failed to suppress crypt hyperproliferation in the colon (fig. S2C-D).

**Fig. 2.**
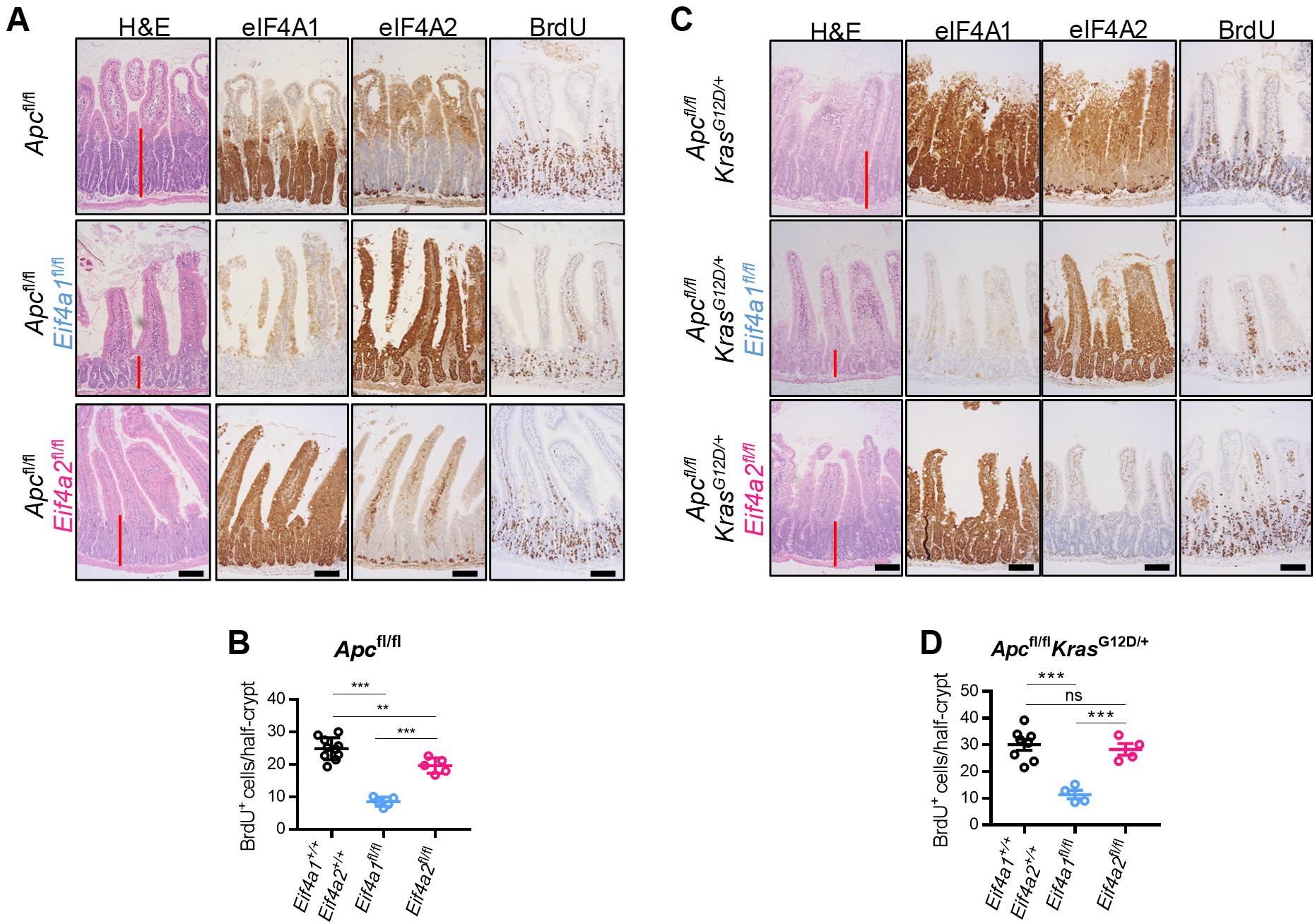
eIF4A1, but not eIF4A2, is essential for Wnt-driven hyperproliferation in the small intestine. **(A)** Representative micrographs of small-intestinal epithelial sections from *VillinCreER^T2^ Apc*^fl/fl^ (*Apc*^fl/fl^) mice harbouring WT eIF4A paralogues, or depletion of either eIF4A1 (*Eif4a1*^fl/fl^) or eIF4A2 (*Eif4a2*^fl/fl^), harvested four days post tamoxifen injection and stained for H&E, eIF4A1, eIF4A2, and BrdU. *Apc*^fl/fl^, n=10; *Apc*^fl/fl^ *Eif4a1*^fl/fl^, n=5; *Apc*^fl/fl^ *Eif4a2*^fl/fl^, n=5. Red bars indicate the expanded crypt depth observed upon *Apc* loss. Scale bars, 100 μm. **(B)** Quantification of BrdU incorporation in half-crypts from mice in (A). Data, mean ± SEM. *Apc*^fl/fl^ vs *Apc*^fl/fl^ *Eif4a1*^fl/fl^, p<0.001; *Apc*^fl/fl^ vs *Apc*^fl/fl^ *Eif4a2*^fl/fl^, p=0.0089; *Apc*^fl/fl^ *Eif4a1*^fl/fl^ vs *Apc*^fl/fl^ *Eif4a2*^fl/fl^, p<0.001. **(C)** Representative micrographs of small-intestinal epithelia from *VillinCreER^T2^ Apc*^fl/fl^ *Kras*^G12D/+^ (*Apc*^fl/fl^ *Kras*^G12D/+^) (n=8), *Apc*^fl/fl^ *Kras*^G12D/+^ *Eif4a1*^fl/fl^ (n=4) or *Apc*^fl/fl^ *Kras*^G12D/+^ *Eif4a2*^fl/fl^ (n=4) mice, stained with H&E, eIF4A1, eIF4A2, or BrdU at day 3 post induction. Red bars indicate the crypt depth observed upon *Apc* loss and *Kras*^G12D/+^ activation. Scale bars, 100 μm. **(D)** Quantification of BrdU incorporation in half-crypts from mice in (C). Data, mean ± SEM. *Apc*^fl/fl^ *Kras*^G12D/+^ vs *Apc*^fl/fl^ *Kras*^G12D/+^ *Eif4a1*^fl/fl^, p<0.001; *Apc*^fl/fl^ *Kras*^G12D/+^ vs *Apc*^fl/fl^ *Kras*^G12D/+^ *Eif4a2*^fl/fl^, p=0.83; *Apc*^fl/fl^ *Kras*^G12D/+^ *Eif4a1*^fl/fl^ vs *Apc*^fl/fl^ *Kras*^G12D/+^ *Eif4a2*^fl/fl^, p<0.001. p-values were calculated by one-way ANOVA and Tukey’s multiple comparisons test. ns = not significant, ** <0.01, *** <0.001.

Expression of eIF4A1, but not eIF4A2, was also required for crypt hyperproliferation in *VillinCre^ER-T2^ Apc^fl/fl^ Kras^G12D/+^* (hereafter *Apc^fl/fl^ Kras^G12D/+^*) mice (Fig. 2C-D and fig. S2E-H), which exhibit constitutively active KRAS signalling. These findings have significant relevance to human disease as *KRAS*-mutated CRC patients represent a hard-to-treat population of clinical unmet need (*49, 50*). For example, while mTOR inhibition can block hyperproliferation following the loss of *Apc*, combining *Apc* loss with Kras^G12D^ activation confers resistance to this treatment (*51, 52*).

### eIF4A1 and eIF4A2 have opposing roles in CRC tumourigenesis

As loss of eIF4A1 is able to supress Wnt-driven crypt hyperproliferation and compromise stem cell maintenance, we next tested whether eIF4A1 is required for CRC tumourigenesis. For this, we first employed the *Lgr5Cre^ER-T2^ Apc^fl/fl^* mouse model, which restricts homozygous loss of *Apc* specifically to *Lgr5⁺* intestinal stem cells, leading to tumour outgrowth. Loss of eIF4A1 led to a significant extension of survival in this model (Fig. 3A), demonstrating that eIF4A1 plays an important role in the intestinal tumourigenesis driven by *Apc* loss. Crucially, all tumours that developed in *Lgr5Cre^ER-T2^ Apc^fl/fl^ Eif4a1^fl/fl^* mice expressed eIF4A1 (fig. S3A) which likely arose from unrecombined cells, suggesting that eIF4A1 is essential for tumour formation.

**Fig. 3.**
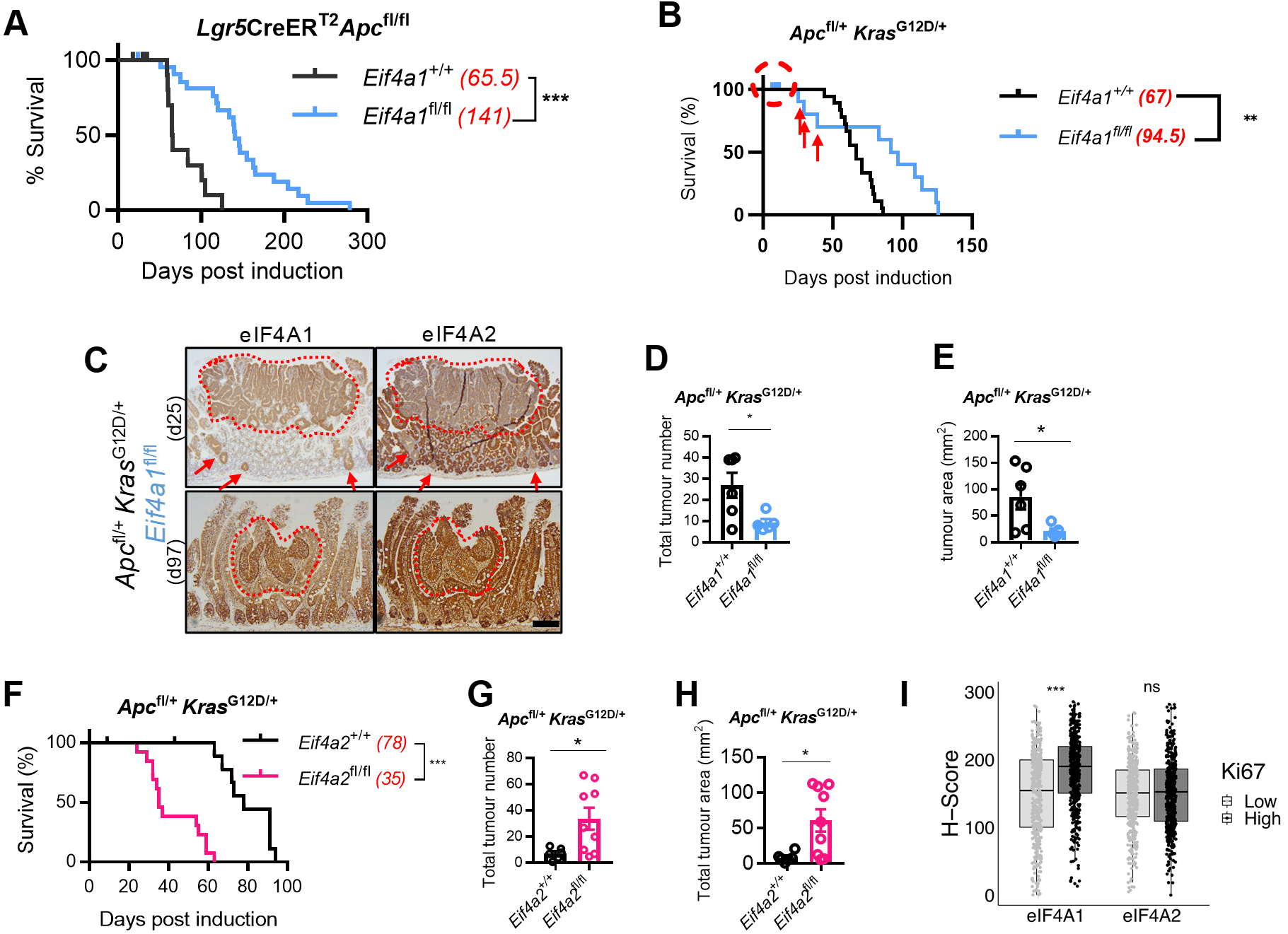
eIF4A1 and eIF4A2 have opposing roles during CRC tumourigenesis. **(A)** Survival curves of *Lgr5Cre^ER-T2^Apc*^fl/fl^ (n=13) and *Lgr5Cre^ER-T2^Apc*^fl/fl^ *Eif4a1*^fl/fl^ (n=23) mice sampled at clinical endpoint. Median survival in days indicated in brackets. Censored mice denoted as tick marks at indicated times post induction; p<0.001. **(B)** Survival curves of *VillinCreER^T2^ Apc*^fl/+^*Kras*^G12D/+^ (*Apc*^fl/+^*Kras*^G12D/+^) (n=18) and *Apc*^fl/+^*Kras*^G12D/+^ *Eif4a1*^fl/fl^ (n=14) mice induced with 80 mg/kg tamoxifen and sampled at clinical endpoint. Median survival in days indicated in brackets. Censored mice denoted as tick marks at indicated times post induction, and early endpoint specimens are indicated with red arrows; p=0.0043. **(C)** Micrographs of representative intestinal tumours (red dotted line) stained for eIF4A1 and eIF4A2 from *Apc*^fl//+^*Kras*^G12D/+^*Eif4a1*^fl/fl^ animals that reached clinical endpoint at either day 25 (top panels) or day 97 (bottom panels) post induction with 80 mg/kg tamoxifen. Early dysplastic crypts positive for eIF4A1 expression are also indicated (red arrows). Scale bar, 100 μm. **(D** and **E)** Total tumour number (G) and total tumour area (H) of *Apc*^fl/+^*Kras*^G12D/+^ (n=6) and *Apc*^fl/+^ *Kras*^G12D/+^*Eif4a1*^fl/fl^ (n=5) mice, induced with 40 mg/kg tamoxifen, sampled around the time corresponding to the median survival of *Apc*^fl/+^*Kras*^G12D/+^ mice (54–61 days). Data, mean ± SEM. (G) p= 0.027; (H) p= 0.037. **(F)** Survival curves of *Apc*^fl/+^*Kras*^G12D/+^ (n=11) and *Apc*^fl/+^*Kras*^G12D/+^*Eif4a2*^fl/fl^ (n=13) mice sampled at clinical endpoint. Median survival in days indicated in brackets. Censored mice denoted as tick marks at indicated times post induction; p<0.001. **(G** and **H)** Total tumour number (J) and total tumour area (K) of *Apc*^fl/+^*Kras*^G12D/+^ (n=6) and *Apc*^fl/+^*Kras*^G12D/+^ *Eif4a2*^fl/fl^ (n=9) mice sampled at day 30 or earlier if the clinical endpoint was reached. Data, mean ± SEM. (J) p= 0.026; (K) p= 0.019. **(I)** Boxplot depicting case-level mean H-scores for eIF4A1 or eIF4A2 expression in human CRC samples stratified by a previously determined clinically relevant Ki67 index threshold of 30% (*79*). Each dot represents the mean H-score, across 1-6 cores, from a unique CRC case (n=900 cases). eIF4A1, p< 0.001; eIF4A2, p= 0.41. p-values for survival data (A, B and F) were obtained with log-rank (Mantel–Cox) tests. p-values for tumour number/area and H-score (D-E, G-I) were calculated with unpaired two-tailed t-tests. ns = not significant, *p <0.05, ** <0.01, *** <0.001.

To investigate this further, we utilised the *VillinCre^ER-T2^ Apc*^fl/+^ *Kras*^G12D/+^ (hereafter *Apc*^fl/+^ *Kras*^G12D/+^) mouse model of CRC development, in which recombination occurs throughout the intestinal epithelium. In this model, the second copy of *Apc* is spontaneously lost over time through a loss-of-heterozygosity event, permitting adenoma outgrowth from the intestinal epithelium. *Apc*^fl/+^ *Kras*^G12D/+^ *Eif4a1*^fl/fl^ mice exhibited significant survival extension with a latency period of roughly 50% longer than that of *Apc*^fl/+^ *Kras*^G12D/+^ control mice (Fig. 3B). Again, at clinical endpoint, all tumours in *Apc*^fl/+^ *Kras*^G12D/+^ *Eif4a1*^fl/fl^ mice expressed eIF4A1 and had low eIF4A2 expression (Fig. 3C), further demonstrating the absolute requirement of eIF4A1 in driving tumourigenesis. Consistent with the homeostasis model, most *Eif4a1*^fl/fl^ mice also harboured intestinal epithelium repopulated by eIF4A1 positive cells (Fig. 3C, d97). However, a few *Apc*^fl/+^ *Kras*^G12D/+^ *Eif4a1*^fl/fl^ mice exhibited early weight loss, with or without tumour formation (appearing as censored or indicated with arrows in Fig. 3B respectively). These mice had retained significant amount of eIF4A1 null cells (Fig. 3C, d25). To circumvent this issue, a lower dose of tamoxifen was used, which also led to a significant extension in survival, with no toxicity (fig. S3B). Taken together, these results suggest that eIF4A1 loss leads to survival extension by delaying tumour formation until the intestine has been repopulated by eIF4A1 expressing clones capable of driving tumourigenesis. Indeed, tumour analyses at 60 days post induction, showed *Apc*^fl/+^ *Kras*^G12D/+^ *Eif4a1*^fl/fl^ mice to have significantly decreased tumour number and reduced tumour volume, compared to of *Apc*^fl/+^ *Kras*^G12D/+^ mice (Fig. 3D-E).

Loss of eIF4A2 in the *Apc*^fl/+^ *Kras*^G12D/+^ model had the opposite effect on CRC development, with *Apc*^fl/+^ *Kras*^G12D/+^ *Eif4a2*^fl/fl^ mice displaying decreased survival, a shorter latency period (Fig. 3F and fig. S3C) and an increased tumour number and burden at 30 days post induction (Fig. 3G-H). Endpoint tumours were negative for eIF4A2 and showed increased eIF4A1 expression (fig. S3D). To assess whether the increase in tumourigenesis, following loss of eIF4A2, was due to the increased expression of eIF4A1, we dampened this expression by crossing with heterozygous *Eif4a1^fl/+^* mice (fig. S3E). This did not rescue the decreased survival (fig. S3F), suggesting that loss of eIF4A2 alone is sufficient to accelerate tumourigenesis in this model.

Collectively, the above results indicate that loss of eIF4A1 suppresses proliferation and impedes tumour formation, while loss of eIF4A2 accelerates tumourigenesis. Consistent with the specific requirement of eIF4A1 to drive crypt hyper-proliferation, all endpoint tumours in *Apc*^fl/+^ *Kras*^G12D/+^ mice were positive for eIF4A1 expression, whereas eIF4A2 expression was detected in a significantly smaller proportion of tumours (fig. S3G-H). To test whether eIF4A1 is specifically required to drive hyper-proliferation also in human disease, we first looked in normal human colonic epithelium, which showed that eIF4A1 is expressed mainly in the Ki67-positive proliferating cells within the crypts, whereas eIF4A2 is expressed in all epithelial cells (fig. S3I). Critically, examination of eIF4A1 and eIF4A2 expression in tumour tissue from CRC patients showed that tumours with the highest rates of proliferation had significantly increased expression of eIF4A1, but not eIF4A2 (Fig. 3I), supporting our finding that specifically eIF4A1 drives tumourigenesis in CRC.

### eIF4A1 is essential for Myc-dependent Wnt signalling

Loss of eIF4A1 in APC deficient mice (Fig. 2A-B) phenocopies loss of Myc (*2*). We therefore hypothesised that oncogenic Wnt signalling requires a balance between the Myc-dependent upregulation of transcriptional targets and the eIF4A1-dependent translation of these mRNAs. To test whether eIF4A1 and Myc impact the same classes of mRNAs, we first performed RNA-seq to determine whether loss of eIF4A1 also suppresses the Myc-dependent subset of Wnt target genes (*2*) at the mRNA level. Loss of eIF4A1, but not eIF4A2, reduced expression of Myc targets deregulated after Apc loss, in both *Apc^fl/fl^* and *Apc^fl/fl^ Kras^G12D/+^*, but not WT, mice (Fig. 4A). Furthermore, loss of eIF4A1 led to reduced nuclear translocation of β-xscatenin in the intestines of *Apc*^fl/fl^ and *Apc*^fl/fl^ *Kras*^G12D*/+*^ mice (Fig. 4B and S3J), consistent with the requirement of eIF4A1 for Wnt-driven hyperproliferation.

**Fig. 4.**
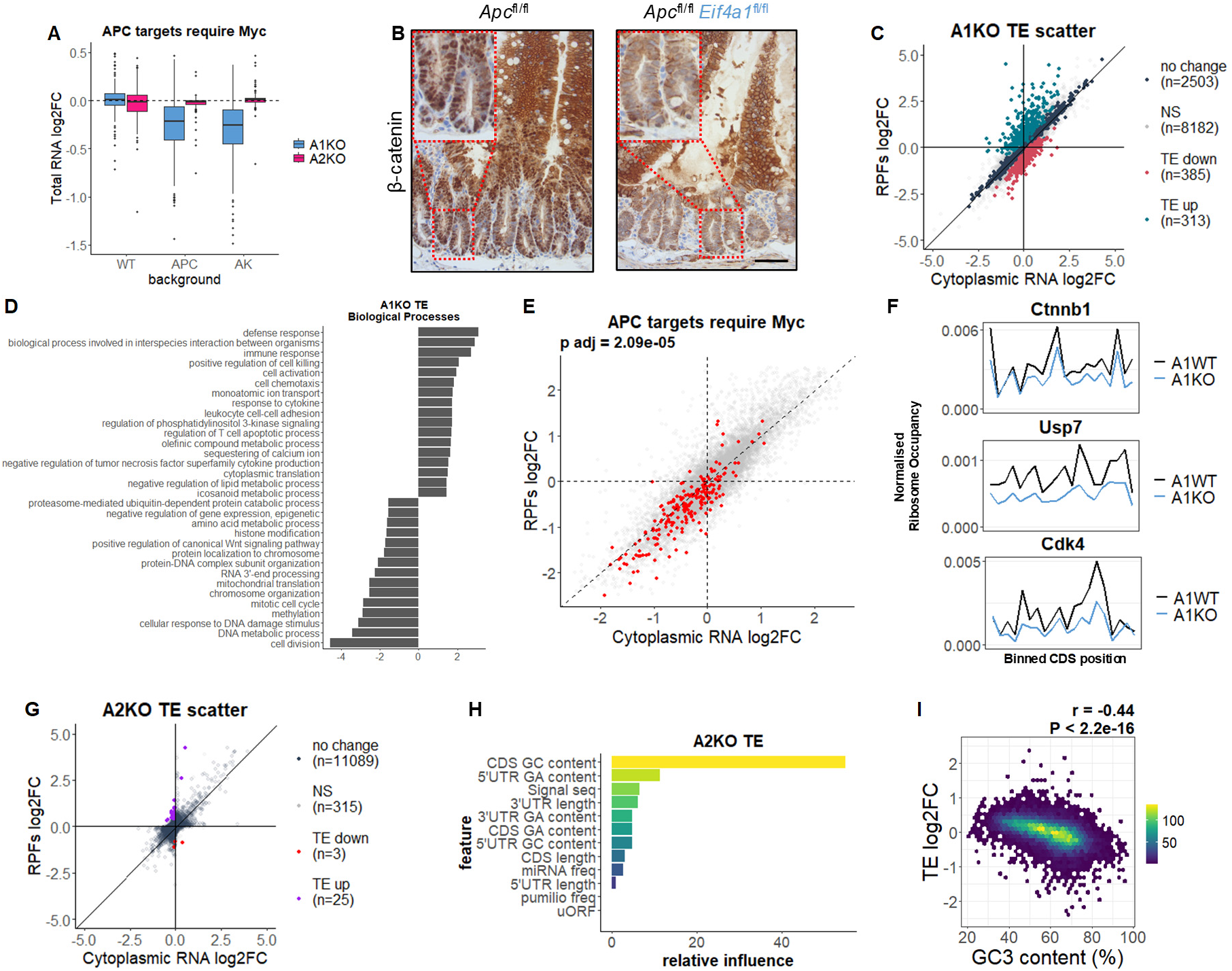
eIF4A1, but not eIF4A2, is required for Wnt-target gene activation following intestinal transformation. **(A)** Boxplot depicting total RNA log2FCs of all mRNAs within the “APC targets require Myc” GSEA gene list (*2*), from RNA-seq of small intestines of *Eif4a1*^fl/fl^ (A1KO) or *Eif4a2*^fl/fl^ (A2KO) mice in either a WT, *Apc*^fl/fl^ (APC), or *Apc*^fl/fl^ *Kras^G12D/+^* (AK) background. n ≥ 4 biological replicates (see methods for exact numbers). A1KO vs A2KO (WT, p= 0.29; APC, p< 0.001; AK, p< 0.001), as calculated with paired two-way t-test. **(B)** β-catenin staining in the small intestine of *Apc*^fl/fl^ and *Apc*^fl/fl^ *Eif4a1*^fl/fl^ mice. Nuclei were counterstained with haematoxylin. Dashed insets show higher magnification of boxed crypt regions. Representative images of n = 3 *Apc*^fl/fl^ and n=4 *Apc*^fl/fl^ *Eif4a1*^fl/fl^ mice. Scale bar, 50 µM. **(C-F)** Ribosome profiling analysis following loss of eIF4A1 in *Apc*^fl/fl^ *Kras*^G12D/+^ small intestines. A1WT = *Apc*^fl/fl^ *Kras*^G12D/+^ *Eif4a1*^+/+^ (n = 4); A1KO = *Apc*^fl/fl^ *Kras*^G12D/+^ *Eif4a1*^fl/fl^ (n = 5). (C) Scatter plot depicting the translational efficiency (TE) changes. Colour scheme depicts the mRNAs identified as translationally down-regulated (padj < 0.1 and TE log2FC < 0), up-regulated (padj < 0.1 and TE log2FC > 0), non-changed (padj ≥ 0.9) or non-significant (NS) (0.1 ≤ padj < 0.9). (D) GSEA using the biological processes as a reference and a ranked list of TE log2FCs as input. Redundant terms have been collapsed using rrvgo, using adjusted p-values as input to calculate scores. Pathways enriched in genes translationally upregulated following eIF4A1 loss have positive scores, and pathways enriched in downregulated genes have negative scores. (E) TE scatter plot, from (C), overlaid with genes from the “APC targets require Myc” GSEA gene list coloured in red. Adjusted p-value was calculated with the fgsea R package. (F) Normalised ribosome occupancy, binned across the length of the CDS, for the indicated Wnt-signalling genes. **(G-I)** Ribosome profiling analysis following loss of eIF4A2 in *Apc*^fl/fl^ *Kras*^G12D/+^ small intestines. A2WT = *Apc*^fl/fl^ *Kras*^G12D/+^ *Eif4a2*^+/+^ (n = 3); A2KO = *Apc*^fl/fl^ *Kras*^G12D/+^ *Eif4a2*^fl/fl^ (n = 4). (G) Scatter plot depicting the translational efficiency (TE) changes. Colour scheme depicts the mRNAs identified as translationally regulated, subset as in (C). (H) Relative influence scores for the feature properties used within a gradient boosting approach, to predict which mRNA features contribute most towards TE log2FCs. **(I)** Density scatter plot comparing GC3 content against TE log2FCs. Colour intensity scale indicates the number of transcripts within each hexagon, which is denoted in the legend. r-and p-values were obtained from a Pearson correlation test.

To further examine whether eIF4A1 promotes the selective translation of specific mRNAs to support Wnt signalling in CRC, we performed ribosome profiling on epithelial extracts from the small intestines of *Apc*^fl/fl^ *Kras*^G12D/+^ mice with and without homozygous loss of either *Eif4a1* or *Eif4a2* (fig. S4-5). We were able to identify 385 mRNAs that had significantly decreased translational efficiency (TE) and 313 mRNAs that increased translationally following loss of eIF4A1 (Fig. 4C).

To determine which processes are regulated by eIF4A1 in these tissues, we performed gene set enrichment analysis (GSEA) on a ranked list of TE log2FCs, using the biological processes (Fig. 4D), Hallmark (fig. S6A) and Curated gene sets (table S1). This identified processes involving cell division as well as Wnt signalling and Myc targets to be translationally repressed following loss of eIF4A1 (Fig. 4D-E and fig. S6). This is therefore consistent with the dependency of oncogenic Wnt signalling on eIF4A1 to drive the translational landscape. Our findings that homeostatic proliferation is unaffected (Fig. 1D) and that Myc-dependent Wnt target genes are unaltered, following loss of eIF4A1 in WT tissue (Fig. 4A), suggest that ligand independent proliferative Wnt signalling requires eIF4A1 activity only in the oncogenic setting. Our data suggest that eIF4A1 is required for the synthesis of proteins at all levels of the oncogenic Wnt-signalling cascade, including β-catenin (encoded by *Ctnnb1)*, upstream components such as the tumour-specific Wnt-activator USP7 (*53, 54*) and downstream Wnt-targets such as CDK4 (*55*) (Fig.3F). *Myc* was repressed in both the RPFs and cytoplasmic RNA, following loss of eIF4A1, which is consistent with *Myc* being a target of both eIF4A1 and oncogenic Wnt signalling.

Among the processes that were translationally upregulated following loss of eIF4A1, we identified an immune signature (Fig. 4D), which may suggest an increase in T-cell recruitment. As activation of Wnt signalling, in response to *Apc* loss, drives immune-cell exclusion in CRC (*56*), we reasoned that perturbation of Wnt signalling, following loss of eIF4A1, could also promote T-cell infiltration back into the crypts. This was supported by CD3/4/8 staining, showing increased T-cell numbers within the crypts following loss of eIF4A1 (fig. S7). Together, these findings are consistent with an essential role for eIF4A1 in Wnt signalling following loss of *Apc* within the gut.

### eIF4A2 regulates translation via a codon-dependent mechanism

In contrast to eIF4A1, loss of eIF4A2 elicited much fewer and smaller changes in gene expression, both at the total RNA and translational level, with only 25 and 3 mRNAs translationally up or down-regulated respectively (Fig. 4G). There was a negative correlation between the changes in TE observed following loss of either eIF4A1 or eIF4A2 (fig. S8A), in agreement with these two paralogues playing opposing roles in translation initiation.

Using GSEA on a ranked list of TE log2FCs, we identified several processes that were altered at the translational level following loss of eIF4A2 (fig. S8B-C). The most striking of these was cytoplasmic translation (fig. S8D), with the translational efficiency of TOP mRNAs (that contain 5′ terminal oligopyrimidine tracts and encode key components of the translational machinery) increasing significantly more than that of non-TOP mRNAs (fig. S8E-F).

To elucidate the molecular mechanisms that explain why loss of these highly related paralogues have such diverse effects on translation, we examined the mRNA feature properties that best explain the observed translational changes (Fig. 4H-I and fig. S9). Interestingly, the supervised machine-learning approach, gradient boosting, identified CDS GC content as the strongest determinant in both datasets, with a relative influence over 60% following loss of eIF4A2 (Fig. 4H and S9A). Due to the triplet nature of codons, GC content within the CDS is mainly driven by the third position of each codon, which is often redundant in determining the amino acid that it encodes, but can affect decoding speed, rendering certain codons as optimal and others non-optimal (*57, 58*). There is a strong negative correlation between GC3 content (GC content at the third position of each codon) and TE log2FC, following loss of eIF4A2 (Fig. 4I). The strength of the correlation is highlighted by comparing to the correlation between TE log2FC following loss of eIF4A1, and either 5’UTR length and GC content (fig. S9 F-G), which are two features thought to define dependence on eIF4A1 activity (*27*). eIF4A2 has previously been shown to repress translation in concert with the CCR4-NOT complex (*23, 25, 26*), which has recently been described as the main sensor of codon optimality (*59–61*). Our data are consistent with eIF4A2 repressing the translation of mRNAs through a codon-dependent mechanism. Indeed, plotting the mean relative synonymous codon usage between mRNAs that are either up or down-regulated specifically in the RPFs (fig. S9M), shows a strong GC/AU split across all codons (fig. S9O). In addition, mRNAs enriched for predicted miRNA and Pumilio binding sites, which provide two alternative mechanisms for recruiting the CCR4-NOT complex, are also translationally up-regulated following loss of eIF4A2 (fig S9P-Q) but not of eIF4A1 (fig. S9K-L).

### eFT226 mimics loss of eIF4A1 in CRC models

Given the opposing effects of eIF4A1 and eIF4A2 in CRC and as all current eIF4A inhibitors target both these proteins (*45–47*), we tested the effects of eIF4A inhibition with eFT226 in our CRC models. eFT226 is a first-in-class inhibitor of eIF4A, which was developed through re-optimisation of rocaglamide-A (RocA) (*62*). *Apc*^fl/fl^ *Kras*^G12D/+^ mice, treated with a single dose of eFT226, exhibited significantly reduced proliferation in intestinal crypts as well as complete loss of stem cell marker expression 24 h post treatment administration (Fig. 5A-D). Furthermore, a single dose of eFT226 was sufficient to ablate stem cells within established tumours, as well as in the underlying crypts, in the *Lgr5Cre^ER-T2^ Apc^fl/fl^*mouse model 24 h post drug treatment (Fig. 5E). eFT226 therefore phenocopies loss of eIF4A1 in these CRC models.

**Fig. 5.**
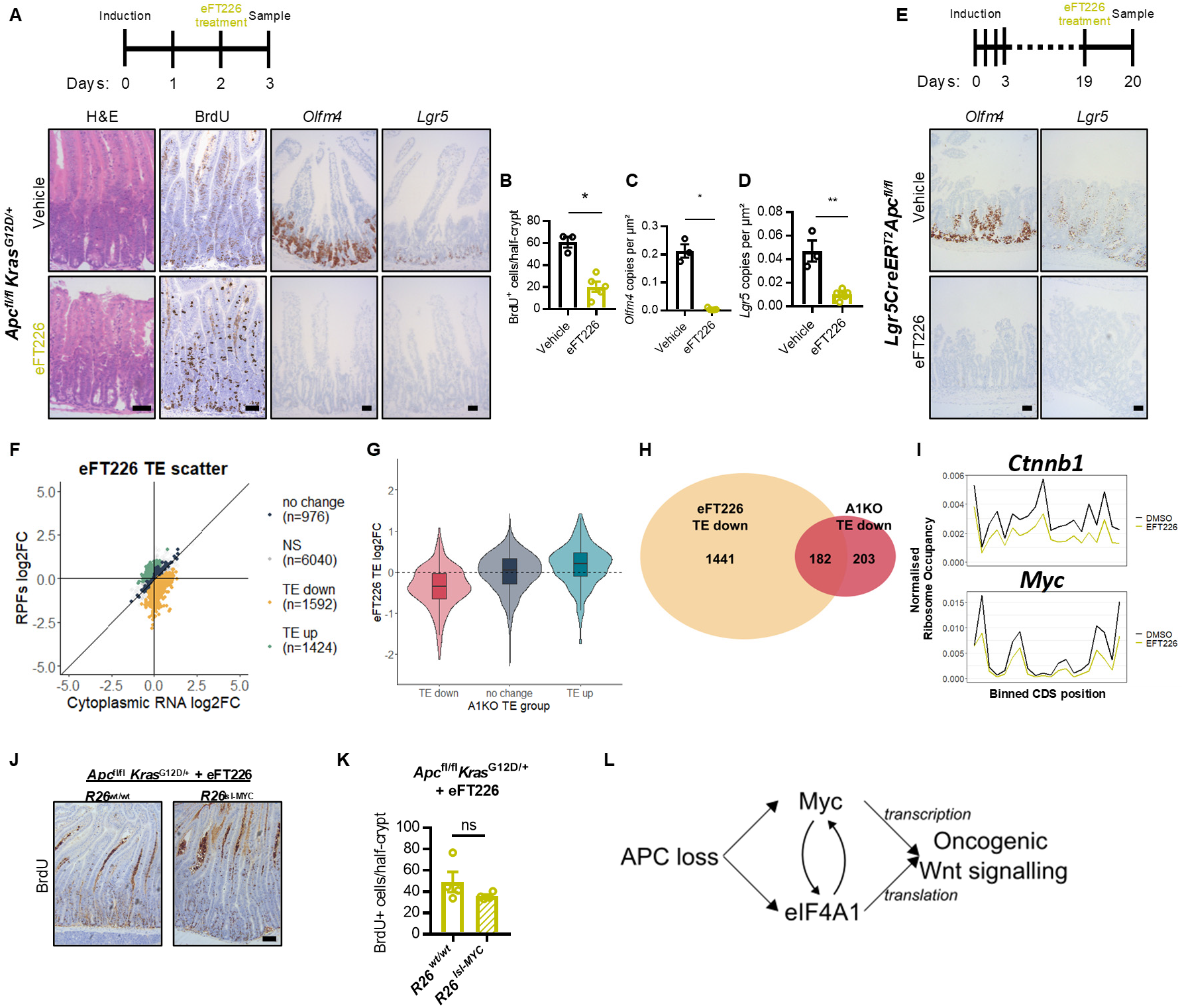
eFT226 mimics loss of eIF4A1 in CRC models. **(A)** Representative micrographs of small-intestinal epithelia from *Apc*^fl/fl^ *Kras*^G12D/+^ mice, treated with eFT226 (n=5) or vehicle (n=3) for 24 h prior to tissue collection at day 3 post induction, and stained with H&E, BrdU (IHC), and for expression of the stem cell markers *Olfm4* and *Lgr5* (ISH). Scale bars, 50 μm. **(B)** Quantification of BrdU⁺ cells in crypts from mice in (A). Data, mean ± SEM shown. Data were statistically assessed by two-tailed Mann–Whitney test; p=0.036. **(C** and **D)** HALO quantification of the expression of the stem cell markers *Olfm4* (C) and *Lgr5* (D) in crypts from mice in (A). Data, mean ± SEM, and statistical significance was assessed by two-tailed Mann–Whitney test in (C), p=0.036, and unpaired two-tailed t-test in (D), p=0.0019. **(E)** Representative micrographs depicting ISH for *Olfm4* and *Lgr5* expression in *Lgr5CreER^T2^ Apc*^fl/fl^ mice, treated with eFT226 or vehicle, 24 h prior to sampling (n=4 mice per group). Scale bars, 50 μm. **(F-I)** Ribosome profiling analysis following 2 h eFT226 treatment in *Apc*^fl/fl^ *Kras*^G12D/+^ small intestines (n = 4 for vehicle and eFT226) (F) Scatter plot depicting the translational efficiency (TE) changes. Colour scheme depicts the mRNAs identified as translationally down-regulated (padj < 0.1 and TE log2FC < 0), up-regulated (padj < 0.1 and TE log2FC > 0), non-changed (padj ≥ 0.9) or non-significant (NS) (0.1 ≤ padj < 0.9). (G) Violin with overlaid boxplots showing TE log2FC, following 2 h eFT226 treatment, for the translationally up-or downregulated groups following loss of eIF4A1 (Fig. 4C). To test for statistical significance, a one-way ANOVA was performed with Tukey’s multiple comparisons to compare all means. padj < 0.001 for all comparisons. (H) Venn diagram depicting the overlap between mRNAs translationally downregulated (“TE down” groups) following eFT226 treatment and loss of eIF4A1. **(I)** Normalised ribosome occupancy, binned across the length of the CDS, for *Ctnnb1* and *Myc*, following 2 h treatment with eFT226 or vehicle. **(J)** Representative micrographs of *Apc*^fl/fl^ *Kras*^G12D/+^ and *Apc*^fl/fl^ *Kras*^G12D/+^*R26^lsl-MYC^*murine intestinal epithelia at three days post induction, treated with eFT226 for 24 h prior to tissue collection (n=4 per group), and stained for BrdU. Scale bar, 50 μm. **(K)** Quantification of BrdU⁺ cells in crypts from mice treated in (J). Data, mean ± SEM. Data were statistically assessed by unpaired two-tailed t-test; p=0.22. **(L)** Proposed model depicting the requirement of both Myc and eIF4A1 for supporting the transcriptional and translational landscape of oncogenic Wnt signalling following loss of *Apc*.

To determine the mechanism whereby eFT226 mimics loss of eIF4A1, we performed ribosome profiling on *Apc*^fl/fl^ *Kras*^G12D/+^ mice, treated with eFT226 or vehicle for 2h (Fig. 5F and S10). The correlations in TE log2FC between eFT226 treatment and either the loss of eIF4A1 or eIF4A2 were poor, when including all mRNAs (fig. S10H-I). However, most mRNAs that were translationally regulated following loss of eIF4A1, were affected in the same direction following eFT226 treatment (Fig. 5G), with 46.6% of eIF4A1-dependent mRNAs also exhibiting translational downregulation following eFT226 treatment (Fig. 5H), including *Ctnnb1* and *Myc* (Fig. 5I). This suggests that eFT226 treatment is mimicking loss of eIF4A1 mainly through the translational repression of an overlapping group of mRNAs. However there is a significantly larger group of mRNAs that are translationally up or down-regulated following eFT226 treatment, but not affected following loss of eIF4A1 (Fig. 5H and S10J). Analysis of the feature properties of these eFT226-sensitive mRNAs suggest that eFT226 is functioning through a distinct gain-of-function mechanism, in which it causes eIF4A to clamp onto purine-rich motifs, thus blocking the scanning ribosome and utilising inhibitory uORFs (fig. S11), consistent with previous data on rocaglamides (*63, 64*). However, both eIF4A1-dependent and eFT226-sensitive mRNAs possess longer 5’UTRs (fig. S9C and S11B), which we believe best explains the strong overlap.

A significant proportion of mRNAs that were translationally upregulated following loss of eIF4A2 were also translationally upregulated following eFT226 treatment, although the majority of these were TOP mRNAs (fig. S12). This is consistent with the possibility that eFT226 can also act to relieve eIF4A2-mediated translational repression. Therefore, while the effect of eFT266 treatment in these CRC model appears to be mainly driven through eIF4A1-dependent mRNAs, this may be distinct in other cancer types.

As *Myc* is translationally repressed following eFT226 treatment (Fig. 5I), it is possible that our data could be explained by the reduction in Myc expression. To test this, we utilised a UTR-less human Myc transgene that has been previously shown to overcome combined mTOR and MNK inhibition in *Apc*^fl/fl^ *Kras*^G12D/+^ mice (*51*). There was no significant difference in the proliferation rates of eFT226 treated *Apc*^fl/fl^ *Kras*^G12D/+^ mice, with or without Myc overexpression (Fig. 5J-K). Together, these findings suggest a model whereby Myc and eIF4A1 are interdependent on each other to support the transcriptional and translational landscape of Wnt-driven CRCs (Fig. 5L).

## Discussion

There is a growing body of work describing the anti-tumourigenic effects of eIF4A inhibitors. However, these drugs target both eIF4A1 and eIF4A2 (*45–47*), with RocA also being recently described to target DDX3X (*65*). We therefore conditionally targeted *Eif4a1* and *Eif4a2* to assess the roles of either paralogue on tumourigenesis. These data demonstrate that the canonical role of eIF4A in promoting translation of oncogenic mRNAs is responsible for driving tumourigenesis in rapidly proliferating cells. We build upon this, by showing that in CRC models, only eIF4A1 has the capacity to support the translational landscape underpinning oncogenic Wnt signalling and single knockouts of eIF4A1 and eIF4A2 have opposite effects on tumourigenesis. Critically, in these models, pharmacological inhibition of eIF4A with eFT226 mimics loss of eIF4A1. Our data suggest that this is due to the translational repression of an overlapping group of mRNAs, despite this occurring through distinct mechanisms.

Our data show that loss of eIF4A1 is sufficient to repress crypt hyper-proliferation in *Apc*^fl/fl^ mice, regardless of *Kras* status, suggesting that these effects are driven predominantly through the essential role of eIF4A1 in promoting the translation of mRNAs that sustain Wnt signalling in the intestine. This is pertinent to the translational relevance of our findings, since more than 80% of CRC patients have loss-of-function mutations in *Apc*, and therefore pharmacological inhibition of eIF4A1 could be highly effective at treating a large population of CRC patients, including those that harbour mutations in *KRAS*.

It is known that *Eif4a1*, but not *Eif4a2*, is a transcriptional target of Myc (*66*), while *Myc* has been shown both here and previously to be a translational target of eIF4A1 (Fig. 5I) (*39, 67*). The cooperativity we observe between Myc and eIF4A1 to transcribe and translate targets of oncogenic Wnt signalling (Fig. 5L) suggests that high expression of one, but not the other, would fail to sustain crypt hyper-proliferation following loss of *Apc* and therefore suggests that inhibiting eIF4A1 could circumvent the need to target Myc directly. Indeed, the importance of balancing the transcriptional and translational landscapes during tumourigenesis has recently been described in bladder cancer, where transcriptional-translational conflict can act as a barrier to tumour progression (*68*).This inter-dependency between Myc and eIF4A1 has also recently been inferred from the observation that eIF4A1-dependency in non-small cell lung cancer is dependent on Myc overexpression (*41*).

We observed a very striking correlation between TE log2FC following loss of eIF4A2 and GC3 content. This suggests that eIF4A2 regulates translation through a codon-dependent mechanism, which is consistent with eIF4A2 acting in concert with the CCR4-NOT complex to sense codon optimality. The increased translation efficiency of mRNAs with AU3 codons (fig. S9N-O), which are enriched for mRNAs associated with cell-cycle terms (*58, 69*), including the GSEA gene sets related to E2F and Myc targets (fig. S8C) as well as TOP mRNAs (fig. S8E-F), could explain why loss of eIF4A2 accelerates CRC tumourigenesis. Conversely, eIF4A2 drives tumorigenesis in hepatocellular carcinoma (HCC), despite also repressing translation in this cancer type (see accompanying manuscript). We submit that this difference reflects distinct routes to tumorigenesis in HCC and CRC. Whilst CRC’s cell-of-origin constitutively resides within a proliferative niche, HCC needs to progress through an extended non-proliferative/senescent-like phase to prime its own tumour-initiation niche.

In summary, we have shown a non-redundant role for eIF4A1 in oncogenic proliferation following loss of *Apc*. Previous studies have suggested that MYC renders many tissues permissive for tumour formation, through increases in translational and proliferative machinery (*70*). Our data argue that a rapid translational response needs to be coordinated to orchestrate these changes. Without this, whilst homeostatic proliferation and differentiation can proceed, the rapid increases in protein synthesis and required translational reprogramming of the proliferative/ribosomal machinery, driven through Wnt-MYC signalling, cannot be sustained. We would thus argue that deregulated/increased translational reprogramming should be considered a prerequisite/hallmark of cancer and not simply a marker of increased proliferation.

## Online Methods

### Mouse studies

All mouse studies were carried out in accordance with UK Home Office regulations (licences 70/8646 and PP3907577), with all protocols and procedures approved by the University of Glasgow Animal Welfare and Ethical Review Board. Mice were genotyped from ear-punch biopsies by Transnetyx. Mice were housed in conventional cages within a pathogen-free facility, maintained at 19–23°C and 55 ± 10% humidity under a 12-h day/night cycle, and allowed access to standard diet and drinking water *ad libitum*. Mice were bred to a C57BL/6J background for at least three generations (n ≥3) before experiments commenced. The *Lgr5CreER^T2^* cohorts were bred for at least seven generations on a C57BL/6J background. The *Apc*^fl/+^ *Kras*^G12D/+^ *Eif4a2*^fl/fl^ ageing experiment was replicated in cohorts bred for ten generations to C57BL/6J mice (fig. S3C). Experiments were performed on mice of both sexes, aged between 2–6 months and weighing ≥ 20 g. Following induction, mice were monitored and weighed minimum of three times per week for signs of ill-health and culled via an approved Schedule 1 method at specified timepoints or once humane endpoint was reached. No formal randomisation was carried out and researchers were not blinded. Animals were censored due to complications with the induction or health issues unrelated to disruption of gut homeostasis or intestinal tumour burden (e.g., epidermal wounds).

Tamoxifen (Sigma, T5648), resuspended in corn oil (Sigma, C8267), was administered by intraperitoneal injection (IP) to induce genetic recombination. The mutant and transgenic alleles used throughout this study were: *VillinCreER^T2^* (*71*), *Lgr5CreER^T2^* (*72*), *Apc*^fl^ (*73*), Lox-Stop-Lox *Kras*^G12D^ (*74*) and *Rosa26-lsl-MYC* (*75*). Generation of the *Eif4a1* and *Eif4a2* floxed alleles is described below.

To induce recombination, tamoxifen was administered via IP at 80 mg/kg for *VillinCreER^T2^* mediated inductions, except in fig. S3B where 40 mg/kg were used. To model the short-term effects of *Apc* loss, *VillinCreER^T2^Apc^fl^*^/fl^ mice were induced with two doses of tamoxifen on consecutive days and sampled 4 days after the first injection, whereas *VillinCreER^T2^ Apc*^fl/fl^ *Kras*^G12D/+^ mice were given a single dose of tamoxifen and sampled 3 days later. To monitor tumourigenesis in ageing cohorts, *VillinCreER^T2^ Apc*^fl/+^ *Kras*^G12D/+^ mice were induced with a single dose of tamoxifen and *Lgr5CreER^T2^ Apc*^fl/fl^ mice were induced with 120 mg/kg tamoxifen on day 0. Mice were then either aged until they presented with clinical symptoms of intestinal tumour burden (hunching, weight loss, and/or anaemia), or were sampled at a pre-specified timepoint. Number and size of tumours were scored macroscopically from methacarn-and/or formalin-fixed intestinal tissue. *Lgr5CreER^T2^ Apc*^fl/fl^ mice that were harvested at day 20 were induced with 120 mg/kg tamoxifen on day 0, followed by three consecutive days with 80 mg/kg tamoxifen.

For eFT226 treatment studies, *Apc*^fl/fl^ *Kras*^G12D/+^ mice were treated with freshly prepared eFT226 administered by a single tail-vein injection of 100 µl 0.25 mg/ml eFT226 solution, prepared in vehicle consisting of 5% DMSO in 10% (w/v) (2-hydroxypropyl)-β-cyclodextrin. eFT226 was provided by Cancer Research Horizons.

### *Eif4a1* and *Eif4a2* floxed allele generation

A conditional allele of *Eif4a1* (Ensembl ID: ENSMUSG00000059796 in Genome Assembly GRCm39) was generated by gene targeting in mouse embryonic stem cells (mESCs). This allele conditionally deletes exons 2–4 (ENSMUSE00001300265, ENSMUSE00001229574 and ENSMUSE00001287896) of the mouse *Eif4a1* gene transcript (Ensembl Transcript ID: Eif4a1-213; ENSMUST00000163666.3) (fig. S13A).

The targeting vector was generated using standard methods and includes an FRT-flanked Neo cassette (*76*). The vector was linearised and transfected into HM1 mESCs (*77*) using electroporation. Cells were selected under G418 (250 µg/ml) and surviving colonies were picked and screened for correct targeting of the *Eif4a1* gene by long-range PCR (using the Roche Expand Long Template PCR System, 11681842001) from within Neo cassette to sequences beyond the ends of the homology arms. Oligo sequences used to screen cells to ensure appropriate targeting of the *Eif4a1* gene were TGGAAACATACTAAATGAACCAGTG and GAGACTAGTGAGACGTGCTACTTCC (2.0kb) for the 5’ side and TCTGAGTAGGTGTCATTCTATTCTGG and CACTGTACATTCCTCAGATCACAAC (2.9 kb) for the 3’ side. The presence of the isolated loxP site was confirmed by PCR with TGACCTAATTACCACCAACTTTACC and CCCCTGACTGATATAAGAACAACAC (166bp).

The conditional allele of the *Eif4a2* gene (Ensembl ID: ENSMUSG00000022884 in Genome Assembly GRCm39) was also generated by gene targeting in mESCs cells. The allele places loxP sites on either side of exons 5 to 10 (ENSMUSE00001211807, ENSMUSE00001217944, ENSMUSE00001310272, ENSMUSE00001291324, ENSMUSE00001263263 and ENSMUSE00001301831) of the mouse *Eif4a2* gene transcript (Ensembl Transcript ID: Eif4a2-201; ENSMUST00000023599.12) (fig. S13B).

The targeting vector was generated, linearised, and transfected as described above, this time with an F3-flanked Neo cassette. Following transfection, surviving colonies were picked and screened. Oligo sequences used to screen cells to ensure appropriate targeting of the *Eif4a2* gene were: AAGTTACAGGCAGGTCAGTCTTCC and CATCGCATTGTCTGAGTAGGTGTC (6.7 kb) for the 5’ side and AGACTAGTGAGACGTGCTACTTCC and TAGACTTGTGATTGGAGCTGGGAG (7.3 kb) for the 3’ side. The presence of the isolated loxP site was confirmed by PCR with GAACAAAGTGGGAAAACTTGTAATG and GCCACTATAACTAAACCAGGGAATC (260 bp).

Following identification of correctly targeted clones for the *Eif4a1* and *Eif4a2* alleles, mouse lines were derived by injection of mESCs cells into C57BL/6J blastocysts according to standard protocols. After breeding of chimeras, germline offspring were identified by coat colour and the presence of the modified allele was confirmed with the 3’ loxP primers described above. Mice were subsequently crossed with a mouse line expressing FLPe (Tg(ACTFLPe)9205Dym) to delete the selectable marker by recombination at the FRT or F3 sites (*78*). Deletion of the selectable marker was confirmed by PCR across the remaining FLP recombinase site. For *Eif4a1*, the oligos used were CGCTTGTCTGTTGTGCAATC and AGATCAAGCGGCTGTAAAGG (494 bp). For *Eif4a2*, the oligo sequences used to confirm deletion of the selectable marker were TAAATTGTCCTAGTCCTGTGAGAGC and TTTTTAAACTTGGGACCTTGTTACC (557 bp).

### Histology and immunohistochemistry

Murine intestinal tissues were fixed in 10% neutral buffered formalin overnight at either room temperature or 4 °C, embedded in paraffin (FFPE), and stained by immunohistochemistry (IHC) and standard histological techniques. IHC was performed on 4 µm FFPE sections which had previously been heated at 60°C for 2 h. Antibodies against the following antigens were applied to tissue sections using a Leica Bond Rx autostainer: BrdU (1:150, BD Biosciences, 347580), eIF4A1 (1:250, Abcam, ab31217), eIF4A2 (1:450, Abcam, ab31218), lysozyme (1:300, DAKO, A0099), β-catenin (1:50, BD Biosciences #610154), CD3 (1:100, Abcam, ab16669), CD4 (1:500, 14-9766-82, eBioscience) and CD8a (1:500, 14-0808-82, eBioscience). All FFPE sections underwent on-board dewaxing (AR9222, Leica) and epitope-retrieval using ER2 retrieval solution (Leica, AR9640) for 20 min at 95 °C. Epitope retrieval for BrdU, eIF4A1, and eIF4A2 was performed using High TRS, while enzymatic retrieval was performed for lysozyme. Sections were rinsed with Leica wash buffer (Leica, AR9590) before peroxidase block was performed using an Intense R kit (Leica, DS9263) for 5 min. Sections were rinsed with wash buffer (Leica, AR9590) before primary antibody application at optimal dilution. The sections were rinsed with wash buffer, incubated with the appropriate secondary antibody, rinsed again with wash buffer, and visualized using DAB from the BOND Intense R Detection kit (Leica, DS9263).

Staining for AB/PAS was performed manually on FFPE sections that were dewaxed and rehydrated through xylene and a graded ethanol series before washing in water. Rehydrated slides were stained for 10 min in Alcian blue solution before rinsing in tap water. Sections were placed in 0.5% periodic acid (Leica, 3803812) for 7 min before washing in tap water and transferred to Schiff’s reagent (CellPath, H5265-500) for 20 min. The sections were washed in tap water to terminate the reaction.

To complete the IHC and AbPAS staining, sections were rinsed in tap water, dehydrated through graded ethanols and placed in xylene. The stained sections were coverslipped in xylene using DPX mountant (CellPath, SEA-1300-00A).

*In situ* hybridisation (ISH) for *Lgr5* (Advanced Cell Diagnostics, 31278) and *Olfm4* (Advanced Cell Diagnostics, 311838) was performed using RNAscope 2.5 LSx reagent kit – brown (Advanced Cell Diagnostics, 322700) on a Bond Rx autostainer (Leica), in accordance with manufacturer’s protocol (Advanced Cell Diagnostics).

To examine cell proliferation, mice were administered 250 μl BrdU (Amersham Biosciences, RPN201) by IP no more than 2 h before sampling. For BrdU-scoring in short-term models, intestinal tissue was fixed in a methacarn solution of methanol (Sigma, 32212), chloroform (Thermo Fisher Scientific, C4960/PB17) and acetic acid (Sigma, 695092) mixed in a ratio of 4:2:1, respectively. Samples were transferred to formalin after 24 h and embedded in paraffin wax. A minimum of 25 half-crypts were scored across intestinal or colonic tissue per biological replicate, and the average number of BrdU⁺ cells per half-crypt per mouse was calculated. A minimum of three mice were analysed per genotype.

### Stain quantification using Halo® software

eIF4A1 and eIF4A2 protein expression levels were quantified using HALO^®^ image analysis software and the Cytonuclear module v1.6 trained to detect intestinal epithelial cells and quantify the optical density of cellular staining. Crypt and villus areas were manually outlined prior to analysis. Analysis was performed in triplicate, with three random areas across three different cross-sections of an intestinal tissue “Swiss roll” analysed per mouse for at least three biological replicates. Staining for eIF4A1 and eIF4A2 was then categorised as negative (0), weak (1+), moderate (2+), or strong (3+), and the Histo-score (H-score) was calculated according to the formula: H-score *=* [1 × (% cells 1+) + 2 × (% cells 2+) + 3 × (% cells 3+)]. Expression of eIF4A1 in the villi was also quantified using Halo^®^. For this, total villus height and villus height stained by the eIF4A1 IHC (from the top of the crypt) were measured. The expanse of eIF4A1 staining is shown as a percentage of total villus height.

### Multiplex immunofluorescence staining of eIF4A paralogues in normal human colon

A multiplex assay consisting of 4 markers was applied to a 4µm thick FFPE section of normal human colon on the Ventana Discovery Ultra autostainer (Roche Tissue Diagnostics, RUO Discovery Universal V21.00.0019). Discovery CC1 (Roche Tissue Diagnostics, 950-123) was applied to the section for 32 minutes at 95°C for antigen retrieval. The antibodies were applied as followed:

Pan-Cytokeratin (AE1/AE3) mouse monoclonal antibody (Leica Biosystems AE1/AE3-601-L-CE) 1:250, followed by Discovery OmniMap anti-Ms HRP (Roche Tissue Diagnostics, 05269652001) detected by Opal 620 (Akoya Bioscience, FP1495001KT) at 1:100, Ki67 (30–9), (Roche Tissue Diagnostics, 790-4286), followed by Discovery OmniMap anti-Rb HRP (Roche Tissue Diagnostics, 760-4311), detected by Opal 570 (Akoya Bioscience, FP1488001KT) at 1:200, anti-eIF4A1 rabbit polyclonal antibody (Abcam, ab31217) 1:50, followed by Discovery UltraMap anti-Rb HRP (RUO) (Roche Tissue Diagnostics, 760-4315), detected by Opal 520 (Akoya Bioscience, FP1487001KT) at 1:500 and finally anti-eIF4A2 rabbit polyclonal antibody (Abcam, ab31218) 1:200, followed by DISCOVERY OmniMap anti-Rb HRP (Roche Tissue Diagnostics, 760-4311), detected by Opal 690 (Akoya Bioscience, FP1497001KT) at 1:300. DAPI was applied as a nuclear counterstain. Whole slide images were acquired at 20x magnification using Phenoimager (Akoya Bioscience, V1.0.13) and unmixed using Phenochart (Akoya Bioscience, V1.1).

### Immunohistochemical quantification of eIF4A paralogues in human CRC samples

Two chromogenic duplex IHC assays were used to quantify the staining of the eIF4A paralogues in 18 CRC tissue microarrays (TMAs) on the Ventana DISCOVERY ULTRA autostainer (Roche Tissue Diagnostics; RUO DISCOVERY Universal procedure, V21.00.0019). Antigen retrieval was performed by treating 4-µm thick FFPE sections with DISCOVERY CC1 solution (Roche Tissue Diagnostics, 950-123) for 32 min at 95 °C. In the first duplex assay, sections were incubated with anti-eIF4A2 antibody (Abcam, ab31218) for 1 h at 1:100, followed by DISCOVERY UltraMap anti-rabbit HRP for 20 min (Roche Tissue Diagnostics, 760-4315), and visualised with the DISCOVERY Purple chromogen (Roche Tissue Diagnostics, 760-229). In the second staining sequence, sections were incubated with anti-cytokeratin antibody (Leica Biosystems, AE1/AE3-601-L-CE) for 28 min at 1:250, followed by DISCOVERY UltraMap anti-mouse AP for 12 min (Roche Tissue Diagnostics, 760-4312), and visualised with the DISCOVERY Yellow chromogen (Roche Tissue Diagnostics, 760-239). For the second duplex assay, sections were incubated with anti-eIF4A1 antibody (Abcam, ab31217) for 1 h at 1:50, followed by incubation with the DISCOVERY UltraMap anti-Rabbit HRP for 16 min (Roche Tissue Diagnostics 760-4315) and detection with the DISCOVERY Purple chromogen (Roche Tissue Diagnostics, 760-229); the second staining sequence detected cytokeratin as described above. Nuclei were stained with haematoxylin and whole-slide images were acquired with the Leica Aperio AT2 digital slide scanner at 40× magnification.

A Visiopharm deep-learning classifier was trained to segment 0.6mm TMA cores into background, tumour, and stroma regions-of-interest using red, green, and blue features at 10× magnification. A further deep-learning classifier was trained to segment tissue into nuclei and background labels using red, green, and blue features at 20× magnification on annotated CRC duplex images. Nuclear labels were refined using post-processing steps to remove objects smaller than 4 µm^2^, fill holes, and separate objects, and a cytoplasmic label was created by dilating nuclei by a distance of 25 µm. A cytoplasmic *H*-score was generated for eIF4A1 and eIF4A2 staining intensity in tumour regions, by manually binning purple stain intensity into 0, 1+, 2+, and 3+ *H*-score bins. Tumour area and percentage-positive tumour area were also calculated.

Case level eIF4A1 and eIF4A2 *H*-scores were calculated as the mean *H*-score from each TMA core for that case. Similarly, the Ki67 index was calculated as the mean Ki67 index (% Ki67⁺ cells) from each TMA core for that case. Only cases that had eIF4A1, eIF4A2, and Ki67 data were selected for further analysis. This resulted in 900 cases with 1-6 cores per case, totalling 2283 cores for eIF4A1 and 2147 cores for eIF4A2. Cases were subset into Ki67-high and Ki67-low using a previously determined clinically relevant Ki67 index threshold of 30% (*79*). Statistical analysis was performed using R (version 4.2.2) and Tidyverse packages (version 2.0.0).

### RNA-seq

Whole small-intestinal tissue was homogenised using CK14 Precellys tubes, followed by on-column DNAse digestion and RNA isolation using RNAeasy kits (QIAGEN #74104), according to the manufacturer’s protocol. RNA concentration was quantified using a NanoDrop 2000c spectrophotometer (Thermo Fisher Scientific) and integrity was checked using an Agilent 220 TapeStation with RNA ScreenTape. Libraries were prepared with the TruSeq RNA sample prep kit v2 (Illumina) and sequenced on an Illumina NextSeq500 instrument, using the High Output 75 cycles kit (2×36 cycles, paired-end reads, single index). *VillinCreER^T2^* (WT, n=5), *Eif4a1*^fl/fl^ (n=6), *Eif4a2*^fl/fl^ (n=4), *Apc*^fl/fl^ (n=4), *Apc*^fl/fl^ *Eif4a1*^fl/fl^ (n=4), *Apc*^fl/fl^ *Eif4a2*^fl/fl^ (n=4), *Apc*^fl/fl^ *Kras*^G12D/+^ (n=4), *Apc*^fl/fl^ *Kras*^G12D/+^ *Eif4a1*^fl/fl^ (n=4) or *Apc*^fl/fl^ *Kras*^G12D/+^ *Eif4a2*^fl/fl^ (n=4).

### Bioinformatic processing of RNA-seq data

Raw FASTQ files were uploaded onto the Gene Expression Omnibus (GEO) database under the accession number GSE240283. Raw sequence quality was assessed using FastQC version 0.11.8, then sequences were trimmed to remove adaptor sequences and low-quality base calls, defined as those with a Phred score < 20, using Trim Galore version 0.6.4. Trimmed sequences were aligned to mouse genome build GRCm38.98, using HISAT2 version 2.1.0, and raw counts per gene were determined using FeatureCounts version 1.6.4. DESeq2 was used for differential expression analysis to calculate log2 fold-changes (log2FCs). These analyses were carried out within each background separately, by comparing single eIF4A paralogue knock-outs to WT. The “APC targets require Myc” mouse gene list (systematic name MM737) was obtained from the Curated GSEA reference (*2*).

### Statistical analyses

All statistical analyses were performed with either GraphPad Prism V7.04 or R version 4.3.1. All details, including the statistical test used to analyse the data and biological replicate numbers for each experiment, are given in the respective figure legend or methods. No statistical method was used to pre-determine the sample sizes. p ≤ 0.05 was considered significant in all cases except for differential expression analyses, in which an adjusted p-value < 0.1 was used, which is default for DESeq2. Data were assessed for normality by D’Agostino–Pearson omnibus K2 test (sample sizes ≥8), or Shapiro–Wilk test (sample sizes <8). Parametric data were assessed by two-tailed t-test, while nonparametric data were compared using two-tailed Mann–Whitney test. Multiple group comparisons were performed with one-way ANOVA. Statistical comparisons of survival data, plotted as Kaplan–Meier curves, were performed using the log-rank (Mantel–Cox) test. Within all boxplots, the horizontal black line inside the box represents the median. Box limits indicate the 25th and 75th centiles. Whiskers represent the maximum and minimum of non-outlier values and extend 1.5× the interquartile range.

### Ribosome profiling

Intestinal crypts were isolated from the proximal part of the small intestine following removal of the villi. To achieve this, intestines were flushed with PBS, cut lengthwise, and incubated at 37°C for 7 min with regular agitation in RPMI 1640 medium (Thermo Fisher Scientific, 21875059) containing 10 mM EDTA and 200 μg/ml cycloheximide. The tissue was then transferred to ice-cold PBS with 10 mM EDTA and 200 μg/mL cycloheximide for 7 min with regular agitation. Tissue remnants were discarded. Crypts were pelleted by centrifugation at 350 *g* for 5 min at 4 °C, snap-frozen in liquid N_2_, and stored at -80 °C.

Pellets were moved from -80 °C to -20°C, roughly 1 h before lysis and then placed on ice for 10 min, prior to lysis with 500 µl ice-cold lysis buffer (15 mM Tris-HCl pH 7.5, 15 mM MgCl_2_, 300 mM NaCl, 1% Triton X-100, 0.05% Tween 20, 2% n-Dodecyl β Maltoside (Thermo Fisher Scientific, 89903), 0.5 mM DTT, 100 µg/ml cycloheximide, 1× cOmplete EDTA-free Protease Inhibitor Cocktail (Merck, 11836170001), 200 U/ml RiboLock RNase Inhibitor (Thermo Fisher Scientific, EO0381)) on ice for 5 min while triturating through a 21G needle with a 1 ml syringe. Lysates were centrifuged at 4°C for 5 min at 12,000 *g* and 500 µl supernatant was pipetted into a fresh 1.5 ml tube. For cytoplasmic RNA samples, RNA was extracted from 25 µl of lysate with 975 µl TRIzol, as per the manufacturer’s instructions. A 20-µl aliquot of lysate was also added to 5 µl of 5× SDS gel-loading buffer for western blotting to check for depletion of eIF4A1 or eIF4A2 in the knock-out samples. The remaining lysate was digested with 10 µl Ambion RNase I (cloned, 100 U/µl; Thermo Fisher Scientific, AM2295) at 22°C for 15 min, with agitation at 600 rpm in a thermo-mixer. The digestion was stopped with 20 µl SUPERaseIn RNase Inhibitor (20 U/μl; Thermo Fisher Scientific, AM2696) and samples were loaded onto a 10–50% sucrose gradient, containing 15 mM Tris-HCl pH 7.5, 15 mM MgCl_2_, 300 mM NaCl, and 100 µg/ml cycloheximide, prepared using a BioComp gradient station and cooled to 4°C for at least 1 h prior to use. Samples were centrifuged in a Beckman XPN-90 Ultracentrifuge with an SW40Ti rotor at 38,000 rpm for 2 h at 4°C. Fractions of 1 ml were collected with a Biocomp gradient station and Gilson FC 203B fraction collector. Fractions containing the 80S ribosome peak were extracted with acid phenol:chloroform, followed by two chloroform washes. RNA was precipitated with 2 µl glycogen (Roche 10901393001), 300 mM NaOAc pH 5.2, and an equal volume of isopropanol overnight at -20 °C.

RNA was pelleted by centrifugation at 12,000 *g* for 45 min at 4 °C. The supernatant was removed with a pipette, and the pellet was washed twice with 70% ethanol and dissolved in 10 µl RNase-free water. RNA was diluted with 10 µl 2× urea buffer (Thermo Fisher Scientific, LC6876), heated at 80°C for 90 s, placed immediately on ice, and then loaded onto a pre-run 15% TBE-urea gel (Thermo Fisher Scientific, EC68852BOX) and ran at 200 V for 1 h alongside custom 28 nt (AGCGUGUACUCCGAAGAGGAUCCAACGU) and 34 nt (GCAUUAACGCGAACUCGGCCUACAAUAGUGACGU) RNA markers. The gel was stained with 1× SYBR gold (Thermo Fisher Scientific, S11494) and imaged on a Typhoon FLA 7000. An image was printed to size to allow bands, inclusive of 28 nt and exclusive of 34 nt markers, to be cut from the gel, placed into a 1.5 ml RNA low-binding microcentrifuge tube, and crushed with a scalpel. RNA was eluted from the crushed gel pieces in 500 µl extraction buffer (300 mM NaOAc pH 5.2, 1 mM EDTA, 0.25% SDS) overnight at 16°C at 600 rpm in a thermo-mixer. Gel pieces were filtered out with a Costar Spin-X centrifuge tube filter (0.45 µm; Scientific Laboratory Supplies Ltd, 8163) and RNA was precipitated with 2 µl glycogen and 500 µl isopropanol overnight at -20 °C.

Precipitated RNA was again pelleted, washed, and dissolved in 43 µl RNase-free water, as above. RPF RNA was not depleted of rRNA. RNA was heated at 80°C for 90 s and immediately placed on ice before undergoing 5′ phosphorylation and 3′ dephosphorylation with 1 µl T4 PNK (NEB, M0201S), 5 µl 10× PNK buffer, and 1 µl SUPERaseIn RNase Inhibitor (20 U/μl) at 37°C for 35 min, with 5 µl 10 mM ATP added for the final 20 min. RNA was extracted with acid phenol:chloroform and isopropanol-precipitated as above.

Purified RNA was quantified on a qubit with the high-sensitivity RNA kit (Q32852) and run on an Agilent bioanalyser to check for correct length. RNA (10 ng) was input into the Bioscientific Nextflex small RNA v3 kit (NOVA-5132-06), using the alternative step F bead clean-up, 11 PCR cycles, and gel-extraction option.

Cytoplasmic RNA samples were run on an Agilent TapeStation to check RNA integrity. RNA concentration and sample purity were measured on a NanoDrop spectrophotometer. rRNA was depleted from 1 µg cytoplasmic RNA with the RiboCop v2, and sequencing libraries were prepared from the rRNA-depleted RNA with the Corall Total RNA-Seq Library Prep Kit v1 (Lexogen, 095.96) using 13 PCR cycles.

Final cytoplasmic RNA and RPF libraries were quantified on an Agilent TapeStation, with a High Sensitivity D1000 tape (Agilent), and sequenced single-end on an Illumina NextSeq500 instrument with a 75 cycles high-output kit.

### Bioinformatic processing of ribosome-profiling data

Processing of all ribosome-profiling data was carried out using the https://github.com/Bushell-lab/Ribo-seq GitHub package. All scripts used within this manuscript can be found in the fork https://github.com/Joseph-Waldron/Ribo-seq_A1_2_colon_paper. All R scripts were carried out using R version 4.3.1. Basic explanations of what these scripts do is written below.

### RPF reads

Sequencing data were demultiplexed with bcl2fastq. Fastq files from the same samples that were split across different runs were concatenated and QC-checked with fastQC. These raw concatenated files were uploaded to the Gene Expression Omnibus (GEO) database with the accession number: GSE240367. Cutadapt (version 1.18) (*80*) was used to remove adaptors, trim 3’ bases with Phred scores < 20, and discard reads fewer than 30 and more than 50 bases after trimming. Unique molecular indexes (UMIs)—4 nt of random sequence at the start and end of every RPF read—were extracted from the reads and appended to the read name using UMI-tools (version 1.0.1) (*81*). BBmap (version 38.18) was used to remove reads that align to either rRNA or tRNA sequences. This was achieved by aligning the reads to FASTA files containing these sequences, writing a new fastq file containing all unaligned reads. These non-rRNA/tRNA reads were then aligned with BBmap to a filtered protein-coding FASTA (see below), containing the most abundant transcript per gene (calculated from cytoplasmic RNA samples). The resulting BAM files were sorted and indexed with samtools (version 1.9) and deduplicated using UMI-tools using the directional method. The number of reads with the 5’ end at each position of every transcript was then counted. The offset for each read length was determined to be 12 nt for read lengths 29–30 nt and 13 nt for read lengths 31–34 nt. These offsets were applied to collapse all reads into a single frame. Total counts across the entire CDS, excluding the first 20 and last 10 codons, were then summed together and used as input to DESeq2 (*82*).

### Cytoplasmic RNA reads

The paired cytoplasmic RNA samples were processed as for the RPFs above but with the following exceptions: The UMIs were 12 nt at the start of each read only. No maximum read length was set when trimming reads with Cutadapt. Reads were aligned to the filtered protein-coding transcriptome with Bowtie2 (version 2.3.5.1) (*83*), using the parameters recommended for use with RSEM, which are --sensitive -- dpad 0 --gbar 99999999 --mp 1,1 --np 1 --score-min L,0,-0.1. Both gene and isoform level quantification was performed using RSEM (version 1.3.3) (*84*). The isoform quantification was used to determine the most abundant transcript per gene (see below), but differential expression was measured at the gene level with DESeq2 (*82*).

### Filtering protein-coding FASTA

The gencode.vM27.pc_transcripts.fa file was downloaded from https://www.gencodegenes.org/mouse/release_M27.html and filtered to include only transcripts that had been manually annotated by HAVANA and that have a 5’UTR, a 3’UTR, a CDS equally divisible by 3, an nUG start codon, and a stop codon. All PAR_Y transcripts were also removed. The cytoplasmic RNA-seq data was then aligned to this filtered FASTA and the most abundant transcript per gene was determined, based on the RSEM output. The RPF reads were then aligned to a FASTA containing only the most abundant transcript for each gene.

### Differential expression analysis

DESeq2 (*82*) was used to test for differential expression in either the RPF or cytoplasmic RNA samples separately, or to test whether the log2FC change in RPFs differed significantly from the log2FC in cytoplasmic RNA, as described previously (*85*), using thresholds described in the figure legends.

### Gene set enrichment analysis

For GSEA (*86*), the R package fgsea (*87*) was used. A ranked list of TE log2FCs was used as input with the mouse-specific Biological Processes, Hallmarks, and Curated reference lists. All output are available in table S1. The rrvgo package (*88*) was used to collapse redundant terms for the biological processes. A threshold of 0.7 was used and -log10(padj values) were used as scores. The mean score within each parent term was extracted and for the terms associated with translational repression, these values were multiplied by -1.

### Single transcript meta plots

For each transcript within a sample, the counts per million (CPM) at every position within that transcript were normalised to the transcripts per million (TPM) value for that transcript for that sample. These normalised CPM values were then subset into 20 equal sized bins, across the length of the CDS, and then averaged within each bin. These were then averaged across each replicate within each sample.

### Relative synonymous codon usage

The relative synonymous codon usage (RSCU) for every codon, across all CDSs, was calculated using the SeqinR R package. The RSCU value reflects codon usage bias and is the frequency of a particular codon relative to the mean frequency of all possible codons that encode each amino acid. It is centred at 1, therefore, if a codon has an RSCU value less than 1, it is used less frequently than expected by chance and *vice versa*. The mean RSCU value for every codon across all CDSs, within each group of transcripts, was plotted and colour-coded by the identity of the wobble nucleotide.

### Feature properties and gradient boosting

The feature properties of all transcripts within the filtered protein-coding transcriptome (see above) were determined using the SeqinR R package. Predicted miRNA-binding sites were extracted from the GSEA M3 subcollection of miRDB microRNA targets, which collects targets obtained from the MirTarget algorithm (*89*) and high confidence predictions (>80) from miRDB v6.0 (*90*). Pumilio-binding sites were determined as UGUANAUA (corresponding to the Pumilio response element consensus sequence) within the 3’UTR (*91*). Transcripts containing upstream open reading frames (uORFs) were determined based on annotation from (*92*), taking those transcripts that had an uORF identified in at least two studies. Transcripts containing a signal sequence were predicted using SignalP 6.0 (*93*).

Gradient boosting was performed using the gbm R package (*94*), using TE log2FC values for all transcripts, assuming a Gaussian distribution, with the following optimised parameters for each dataset:

A1KO: n.trees = 765, interaction.depth = 5, shrinkage = 0.01, cv.folds=5, n.minobsinnode = 10, bag.fraction = 0.75.

A2KO: n.trees = 4988, interaction.depth = 5, shrinkage = 0.01, cv.folds=5, n.minobsinnode = 5, bag.fraction = 0.75.

eFT226: n.trees = 1066, interaction.depth = 5, shrinkage = 0.01, cv.folds=5, n.minobsinnode = 5, bag.fraction = 0.75.

### GA tetramer enrichment

The positional enrichment of GA tetramers was performed as previously described (*23*). The percentage of transcripts, containing one of the eight most-enriched tetramers (AAGA|AGAA|GAAA|GAGA|AGAG|GGAA|AAAA|GAAG) from a previous eIF4A Bind-n-Seq experiment (*63*), was plotted at every described position.

## Acknowledgments

We would like to thank Cancer Research Horizons for the supply of eFT226 and Neil Jones for his intellectual input. We would like to thank Natalia Sphyris and Catherine Winchester for critical reading of the manuscript. The CRUK Scotland Institute acknowledges the support of NHS Research Scotland (NRS) Greater Glasgow and Clyde Biorepository. We acknowledge Silvia Martinelli and Rachel Pennie for their support with human histology.

## Funding

CRUK core funding to the CRUK Scotland Institute (A31287)

CRUK core funding to the CRUK Glasgow Centre (A25142)

CRUK core funding to the CRUK Scotland Centre (CTRQQR-2021\100006)

CRUK core funding to MB lab (A29252) (JAW, JM, SLG)

CRUK core funding to OJS lab (A21139 and DRCQQR-May21\100002) (RCLS, GK, JRPK, CA, NV, ADC, RAR, KG)

Mazumdar-Shaw Chair (JLQ lab)

CRUK core funding to JN lab (A28291) (LPF and MM)

SPECIFICANCER Cancer Research Grand Challenge (A29055) (KG)

CSO fellowship TCS/22/02 (KP)

## Author contributions

Conceptualization: OJS, MB

Methodology, Investigation, Visualization and Supervision: RCLS and GK performed all mouse experiments, under the supervision of JRPK and OJS, with support from CA, NV, ADC, RAR, LPF, MM, JN and SBC. JAW and JM carried out ribosome-profiling, with support from SLG, under the supervision of MB. JAW carried out ribosome-profiling analysis, with support from SLG, under the supervision of MB. KG and JAW carried out RNA-seq analysis. Generation of floxed *Eif4a1* and *Eif4a2* alleles was carried out by DStr, DSte and FCW. Mouse histology was performed by CN, BC and the CRUK Scotland Institute histology services. TMA was sourced under the governance of NHSGGC Biorepository and curated and constructed by JE, KP and JH. Samples were generated by JH and human histology was carried out by FB under the supervision of LOJ and JLQ. Image analysis of human histology was carried out by LOJ, and data analysis was carried out by IP under the supervision of JLQ.

Funding acquisition: OJS, MB, JLQ, JN

Writing – original draft: JAW

Writing – review & editing: GK, RCLS, JRPK, MB and OJS

## Data and materials availability

RNA-seq data has been deposited to the Gene Expression Omnibus (GEO) database under the accession number GSE240283. Ribosome profiling data has been deposited to GEO under the accession number GSE240367.

