## Supplemental figures for "eIF4A1 is essential for reprogramming the translational landscape of Wnt-driven colorectal cancers"

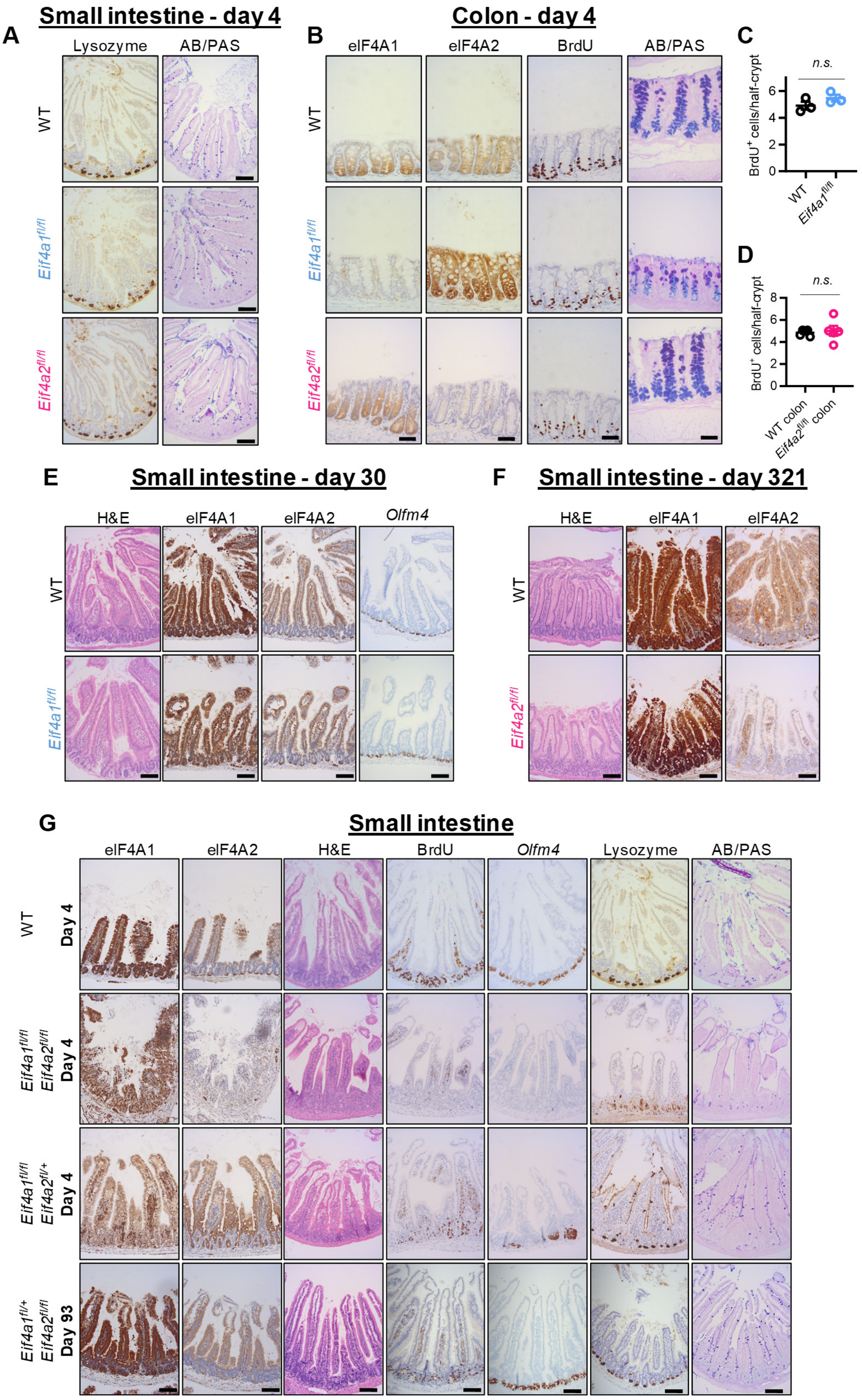


**Fig. S1. (legend on following page)**

**Targeting *Eif4a1* and *Eif4a2* in the WT mouse intestine. (A)** Representative micrographs of murine small-intestinal epithelial sections from *VillinCreER$^{T2}$* (WT, n=5), *VillinCreER$^{T2}$ Eif4a1*$^{fl/fl}$ (*Eif4a1*$^{fl/fl}$, n=5), or *VillinCreER$^{T2}$ Eif4a2*$^{fl/fl}$ (*Eif4a2*$^{fl/fl}$, n=5) mice sampled four days post induction and stained for lysozyme (Paneth cells) and alcian blue/PAS (goblet cells). Scale bar, 100 µm. **(B)** Representative micrographs of colon tissue from mice in (A) stained for eIF4A1, eIF4A2, BrdU, and alcian blue/PAS. Scale bars, 50 µm. **(C** and **D)** Quantification of BrdU$^{+}$ cells following loss of eIF4A1 or eIF4A2 in the colon. Data are presented as mean ± SEM and were statistically assessed by unpaired two-tailed t-tests. (C) WT, n=3; *Eif4a1*$^{fl/fl}$, n=3; p=0.11. (D) WT, n=5; *Eif4a2$^{fl/f}$,* n=5; p=0.74. **(E)** Representative micrographs of H&E, eIF4A1, eIF4A2 and *Olfm4* staining in intestinal tissue from WT (n=3) and *Eif4a1*$^{fl/fl}$ mice (n=2) sampled 30 days post induction. Scale bars, 100 µm. **(F)** Representative micrographs of H&E, eIF4A1, and eIF4A2 staining in intestinal tissue from WT (n=4) and *Eif4a2*$^{fl/fl}$ mice (n=2) sampled 321 days post induction. Scale bars, 100 µm. **(G)** Representative micrographs of small-intestinal epithelia from mice harbouring concurrent deletion of *Eif4a1* and *Eif4a2* (*Eif4a1*$^{fl/fl}$ *Eif4a2*$^{fl/fl}$, n=4), or mice harbouring homozygous loss of *Eif4a1* and single allele deletion of *Eif4a2* (*Eif4a1*$^{fl/fl}$ *Eif4a2$^{fl/+}$*, n=4) four days post induction. Representative micrographs are also shown for a cohort with monoallelic *Eif4a1* expression and homozygous loss of *Eif4a2* (*Eif4a1$^{fl/+}$ Eif4a2*$^{fl/fl}$, n=3), sampled 93 days post induction. Tissue sections were stained for eIF4A1, eIF4A2, H&E, BrdU, *Olfm4* (ISH; stem cells), lysozyme (Paneth cells), and alcian blue /PAS (goblet cells). Scale bars, 100 µm. Please note that the WT control panels are shared with Fig. 1A.

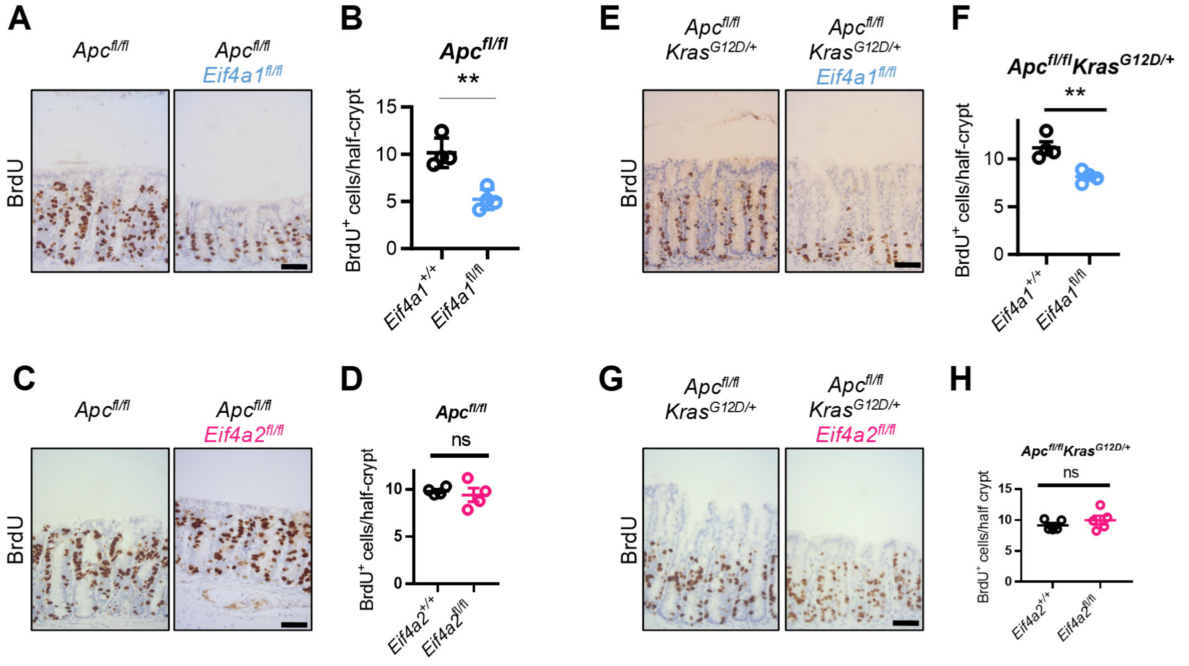


**Fig. S2.**
**eIF4A1, but not eIF4A2, is essential for Wnt-driven hyperproliferation in the colon. (A-D)** BrdU IHC staining and quantification in colons of *VillinCreER*$^{T2}$ *Apc*$^{fl/fl}$ (*Apc*$^{fl/fl}$) mice with and without loss of eIF4A1 (A and B) or eIF4A2 (C and D) at day 4 post induction. n=4 per group. Scale bars, 50 µm. (B) p=0.002, (D) p=0.60 **(E** to **H)** BrdU IHC staining and quantification in colons of *Apc*$^{fl/fl}$*Kras*$^{G12D/+}$ mice with and without loss of eIF4A1 (n=4) (E and F) or eIF4A2 (n=5) (G and H) at day 3 post induction. (F) p=0.0047, (H) p=0.33. Scale bars, 50 µm. Data, mean ± SEM. Data were statistically assessed by unpaired two-tailed t-tests. ns = not significant, ** p<0.01.

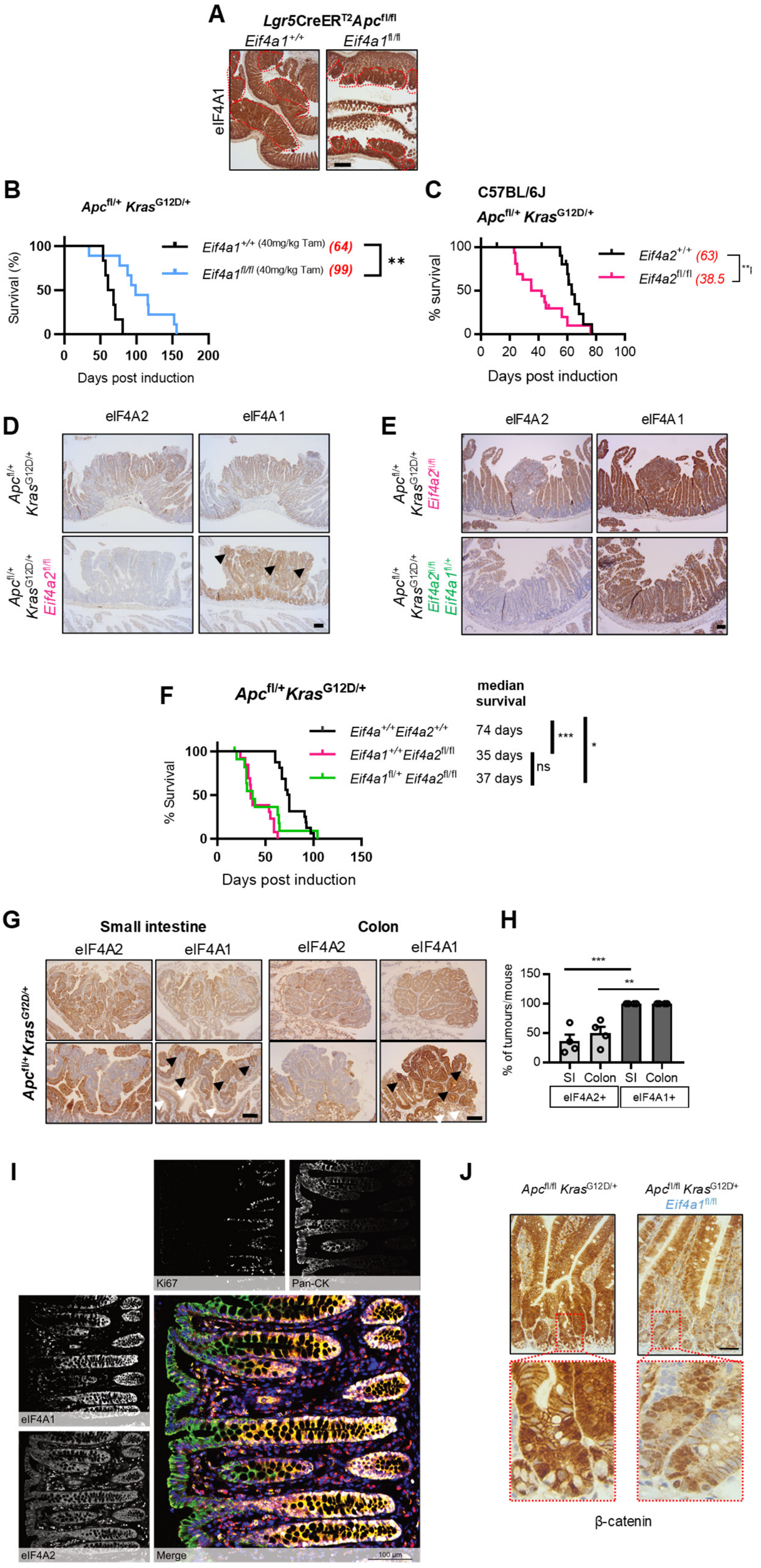

**Fig. S3. (legend on following page)**

**eIF4A1 and eIF4A2 have opposing roles during CRC tumourigenesis. (A)** Representative micrographs of eIF4A1 IHC staining in the intestine of indicated mice collected at clinical endpoint for Fig. 3A. Individual tumours outlined with dotted lines. Scale bar, 400 µm. **(B)** Survival curves of *VillinCreER*$^{T2}$ *Apc*$^{fl/+}$*Kras*$^{G12D/+}$ (*Apc*$^{fl/+}$*Kras*$^{G12D/+}$) (n=6) and *Apc*$^{fl/+}$*Kras*$^{G12D/+}$ *Eif4a1*$^{fl/fl}$ (n=9) mice induced with 40 mg/kg tamoxifen sampled at clinical endpoint. Median survival in days indicated in brackets. p=0.0023 as calculated by log-rank (Mantel–Cox) test. **(C)** Survival curves of *Apc*$^{fl/+}$ *Kras*$^{G12D/+}$ (n=12) and *Apc*$^{fl/+}$ *Kras*$^{G12D/+}$ *Eif4a2*$^{fl/fl}$ (n=16) mice bred to a C57BL/6J background for at least 10 generations and sampled at clinical endpoint. Median survival in days indicated in brackets. p=0.006 as calculated by log-rank (Mantel–Cox) test. Censored mice denoted as tick marks at indicated times post induction. This shows that the data from Fig. 3F could be recapitulated in *Apc*$^{fl/+}$ *Kras*$^{G12D/+}$ *Eif4a2*$^{fl/fl}$ cohorts bred to a C57BL/6J background for at least ten generations. **(D)** Representative micrographs of eIF4A2 and eIF4A1 expression in intestinal tumours of *Apc*$^{fl/+}$ *Kras*$^{G12D/+}$ and *Apc*$^{fl/+}$ *Kras*$^{G12D/+}$ *Eif4a2*$^{fl/fl}$ mice. Black arrow heads point to increased eIF4A1 expression in *Eif4a2*$^{fl/fl}$ tumour epithelia. Scale bar, 100 µm. **(E)** Representative micrographs of eIF4A2 and eIF4A1 expression in intestinal tumours of *Apc*$^{fl/+}$ *Kras*$^{G12D/+}$ *Eif4a2*$^{fl/fl}$ and *Apc*$^{fl/+}$ *Kras*$^{G12D/+}$ *Eif4a1*$^{fl/+}$ *Eif4a2*$^{fl/fl}$ mice at clinical endpoint. Scale bar, 100 µm. **(F)** Survival curves of *Apc*$^{fl/+}$ *Kras*$^{G12D/+}$ (n=16), *Apc*$^{fl/+}$ *Kras*$^{G12D/+}$ *Eif4a2*$^{fl/fl}$ (n=13), and *Apc*$^{fl/+}$ *Kras*$^{G12D/+}$ *Eif4a1*$^{fl/+}$ *Eif4a2*$^{fl/fl}$ (n=13) mice sampled at clinical endpoint. Median survival indicated on the right. p-values obtained by log-rank (Mantel–Cox) tests; *Apc*$^{fl/+}$ *Kras*$^{G12D/+}$ vs *Apc*$^{fl/+}$ *Kras*$^{G12D/+}$ *Eif4a2*$^{fl/fl}$ p<0.001, *Apc*$^{fl/+}$ *Kras*$^{G12D/+}$ vs *Apc*$^{fl/+}$ *Kras*$^{G12D/+}$ *Eif4a1*$^{fl/+}$ *Eif4a2*$^{fl/fl}$ p=0.024, *Apc*$^{fl/+}$ *Kras*$^{G12D/+}$ *Eif4a2*$^{fl/fl}$ vs *Apc*$^{fl/+}$ *Kras*$^{G12D/+}$ *Eif4a1*$^{fl/+}$ *Eif4a2*$^{fl/fl}$ p= 0.30. Please note that the *Eif4a2*$^{fl/fl}$ cohort is the same as presented in Fig. 3F. Censored mice denoted as tick marks at indicated times post induction. **(G)** Representative micrographs of small-intestinal and colonic tumours from *Apc*$^{fl/+}$ *Kras*$^{G12D/+}$ mice, at clinical endpoint, stained for eIF4A1 and eIF4A2 expression. Small-intestinal and colonic tumours show variable expression of eIF4A2 (top left panels, eIF4A2-high; bottom left panels. eIF4A2-low), while eIF4A1 is robustly and uniformly expressed throughout the intestinal tumour epithelium. eIF4A1 expression appears increased in eIF4A2-low tumour epithelia (black arrowheads), while eIF4A1 expression in non-tumour epithelia (white arrowheads) appears lower than in tumours. Scale bars, 100 µm. **(H)** Percentage of tumours positive for eIF4A2 and eIF4A1 (n=4). Data, mean ± SEM and assessed with one-way ANOVA and Tukey’s multiple comparisons test; SI; p=0.0004, colon; p=0.0031. **(I)** Normal human colonic epithelium stained for eIF4A1 (yellow), eIF4A2 (red), Ki67 (white), cytokeratin (pan-CK; green)) and cell nuclei (blue) and visualized using multiplex immunofluorescence. **(J)** β-catenin IHC staining in the small intestine of *Apc*$^{fl/fl}$ *Kras*$^{G12D/+}$ and *Apc*$^{fl/fl}$ *Kras*$^{G12D/+}$ *Eif4a1*$^{fl/fl}$ mice. Nuclei were counterstained with haematoxylin. Lower panels are higher magnification of boxed crypt regions. Representative images of n = 3 biological replicates per group. Scale bar, 50 µM.

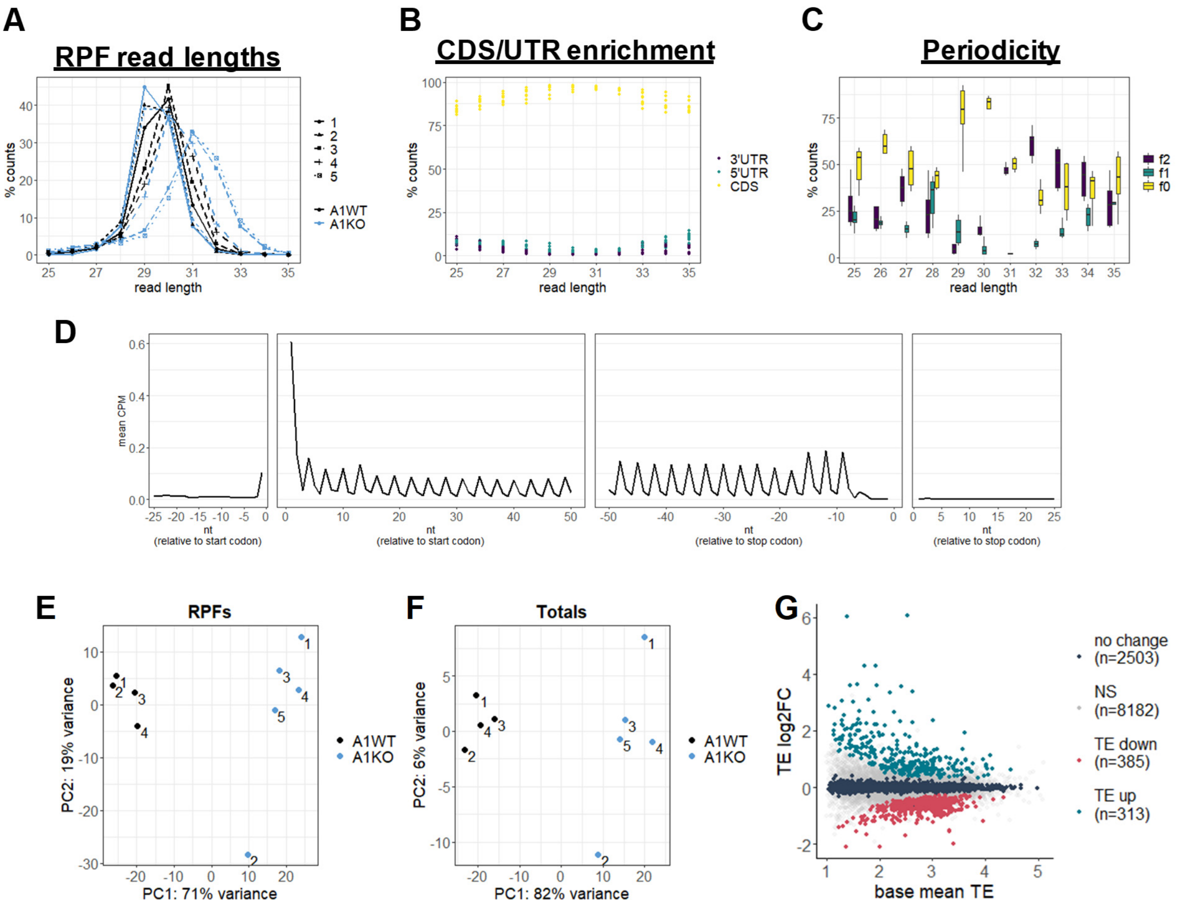


**Fig. S4.**

**Quality control of ribosome-profiling in small intestines following eIF4A1 knockout.**
A1WT = *VillinCreER*$^{T2}$ *Apc*$^{fl/fl}$ *Kras*$^{G12D/+}$ (n=4) and A1KO = *VillinCreER*$^{T2}$ *Apc*$^{fl/fl}$ *Kras*$^{G12D/+}$ *Eif4a1*$^{fl/fl}$ (n=5). **(A)** Ribosome protected fragments (RPF) read length distribution for indicated samples. **(B)** Percentage of protein-coding RPF reads that aligned to either the 5'UTR, CDS, or 3'UTR, separated by read length. **(C)** Boxplot depicting the percentage of RPF reads within each frame of the coding sequence, separated by read length. Read lengths 29–34 were taken forward for downstream analysis as these represented the highest CDS:3'UTR ratio of reads and best periodicity. **(D)** Average counts per million (CPM) at the indicated positions, up and down-stream of the start and stop codons, after applying an offset of 12 nt for read lengths 29–30 and 13 for read lengths 31–34. **(E-F)** Principal component analysis (PCA) plot of all RPF (E) and cytoplasmic RNA (F) samples after batch correction. **(G)** Translational efficiency (TE) MA plot. TE log2FC = RPFs log2FC – Cytoplasmic RNA log2FC. Base mean TE is the mean TE (normalised RPFs - normalised cytoplasmic RNA) across all samples, as calculated with DESeq2.

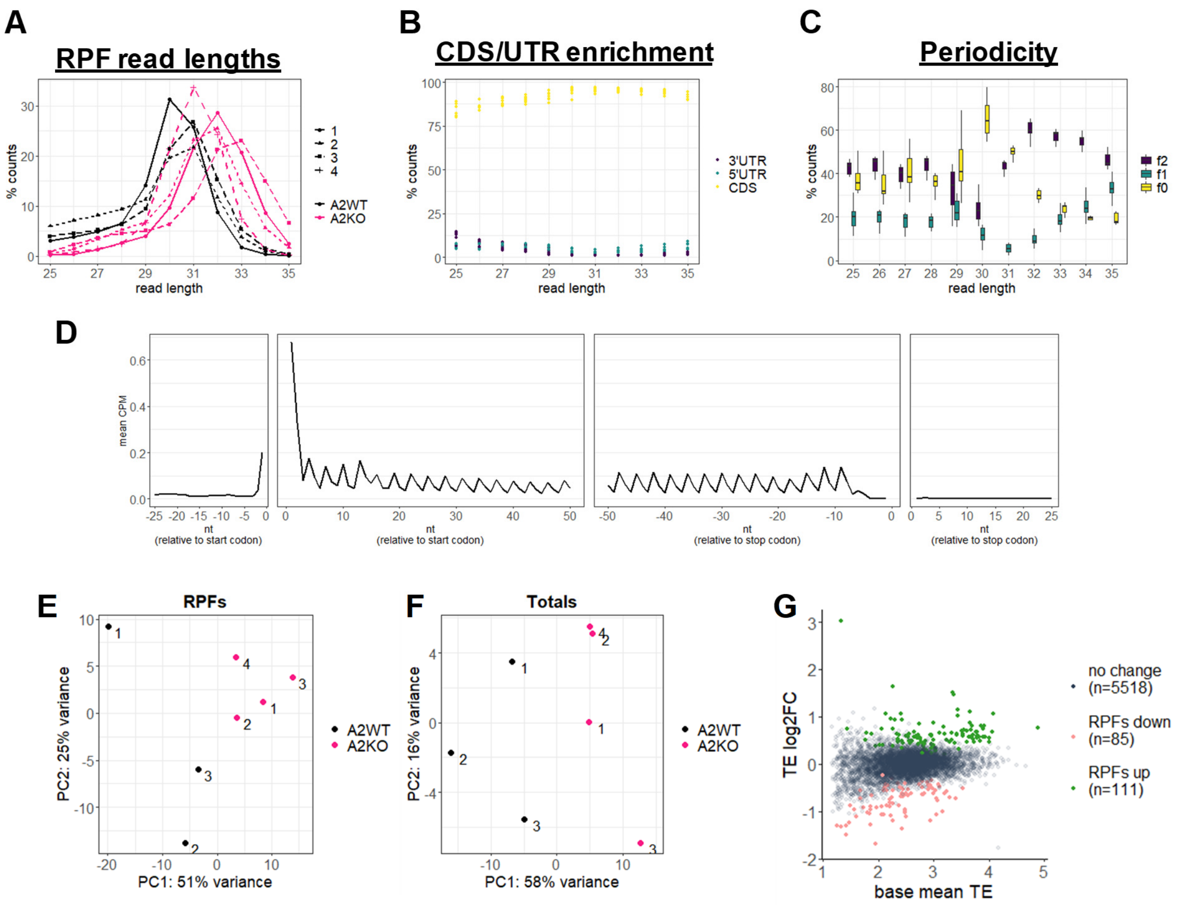


**Fig. S5.**

**Quality control of ribosome-profiling in small intestines following eIF4A2 knockout.**

A2WT = *VillinCreER*$^{T2}$ *Apc*$^{fl/fl}$ *Kras*$^{G12D/+}$ (n=3) and A2KO = *VillinCreER*$^{T2}$ *Apc*$^{fl/fl}$ *Kras*$^{G12D/+}$ *Eif4a2*$^{fl/fl}$ (n=4). **(A)** Ribosome protected fragments (RPF) read length distribution for indicated samples. **(B)** Percentage of protein-coding RPF reads that aligned to either the 5'UTR, CDS, or 3'UTR, separated by read length. **(C)** Boxplot depicting the percentage of RPF reads within each frame of the coding sequence, separated by read length. Read lengths 29–34 were taken forward for downstream analysis as these represented the highest CDS:3'UTR ratio of reads and best periodicity. **(D)** Average counts per million (CPM) at the indicated positions, up and down-stream of the start and stop codons, after applying an offset of 12 nt for read lengths 29–30 and 13 for read lengths 31–34. **(E-F)** Principal component analysis (PCA) plot of all RPF (E) and cytoplasmic RNA (F) samples after batch correction. **(G)** Translational efficiency (TE) MA plot. TE log2FC = RPFs log2FC – Cytoplasmic RNA log2FC. Base mean TE is the mean TE (normalised RPFs - normalised cytoplasmic RNA) across all samples, as calculated with DESeq2.

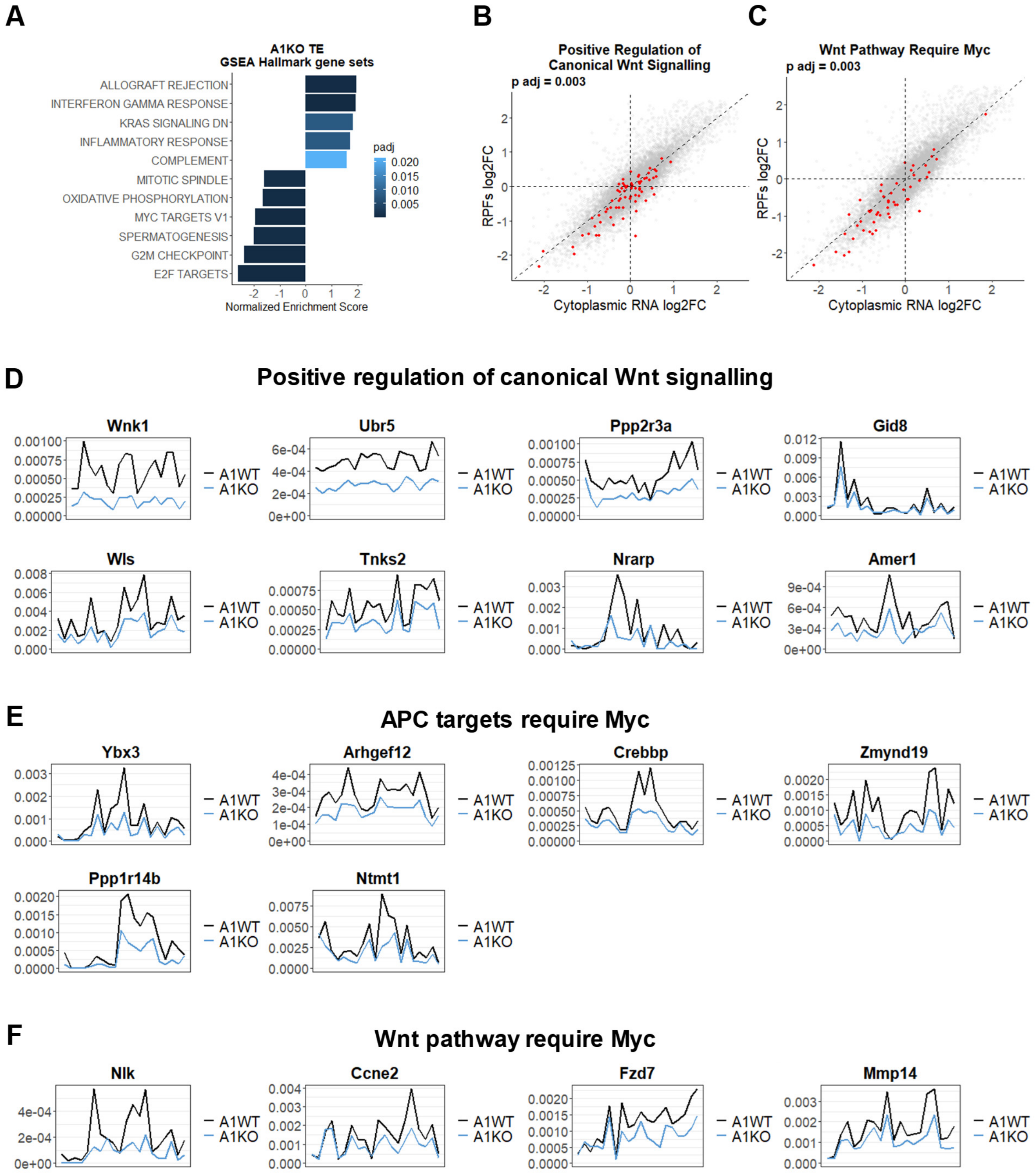


**Fig. S6.**
**eIF4A1 supports the translational landscape of Wnt-driven hyperproliferation.** Ribosome profiling analysis following loss of eIF4A1 in *Apc*$^{fl/fl}$ *Kras*$^{G12D/+}$ small intestines, as in Fig. 4C-F and fig. S4. **(A)** GSEA using the Hallmark gene sets on a ranked list of TE log2FCs, with bar plot showing significantly enriched (padj < 0.05) Hallmark pathways in genes translationally altered following loss of eIF4A1. Pathways are ordered by normalized enrichment score (NES) and coloured by adjusted p-value. Pathways enriched in genes translationally upregulated following eIF4A1 loss have NES >0, and pathways enriched in downregulated genes have NES

<0. **(B-C)** Overlaid TE scatter plots (from Fig. 4C), with genes from the indicated gene list coloured in red. Adjusted p-value was calculated with the fgsea R package. **(D-F)** Meta plots depicting the normalised ribosome occupancy across the length of the CDS for A1KO “TE down” mRNAs from the GSEA gene lists; “Positive regulation of canonical Wnt signalling” (D), “APC targets require Myc” (E) and “Wnt pathway require Myc” (F).

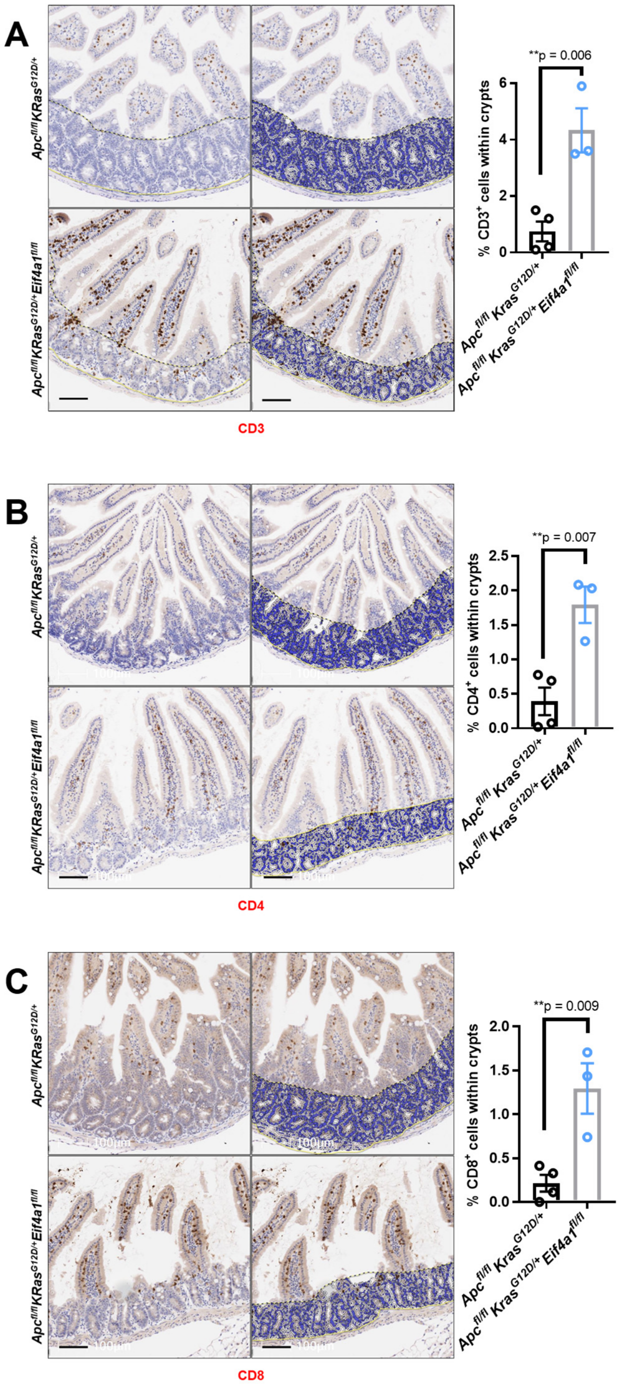


**Fig. S7.**

**Loss of eIF4A1 leads to T-cell infiltration in *Apc*$^{fl/fl}$ *Kras*$^{G12D/+}$ intestines. (A** to **C)** Representative micrographs of *Apc*$^{fl/fl}$ *Kras*$^{G12D/+}$ (n=4) or *Apc*$^{fl/fl}$ *Kras*$^{G12D/+}$ *Eif4a1*$^{fl/fl}$ (n=3) intestines, stained for CD3 (A), CD4 (B), and CD8 (C), three days post induction. Images on the right show HALO rendering and quantification of positively-stained cells, present in intestinal crypts, is shown on the right. Scale bars, 100 μm. Data were statistically assessed by unpaired two-tailed t-tests.

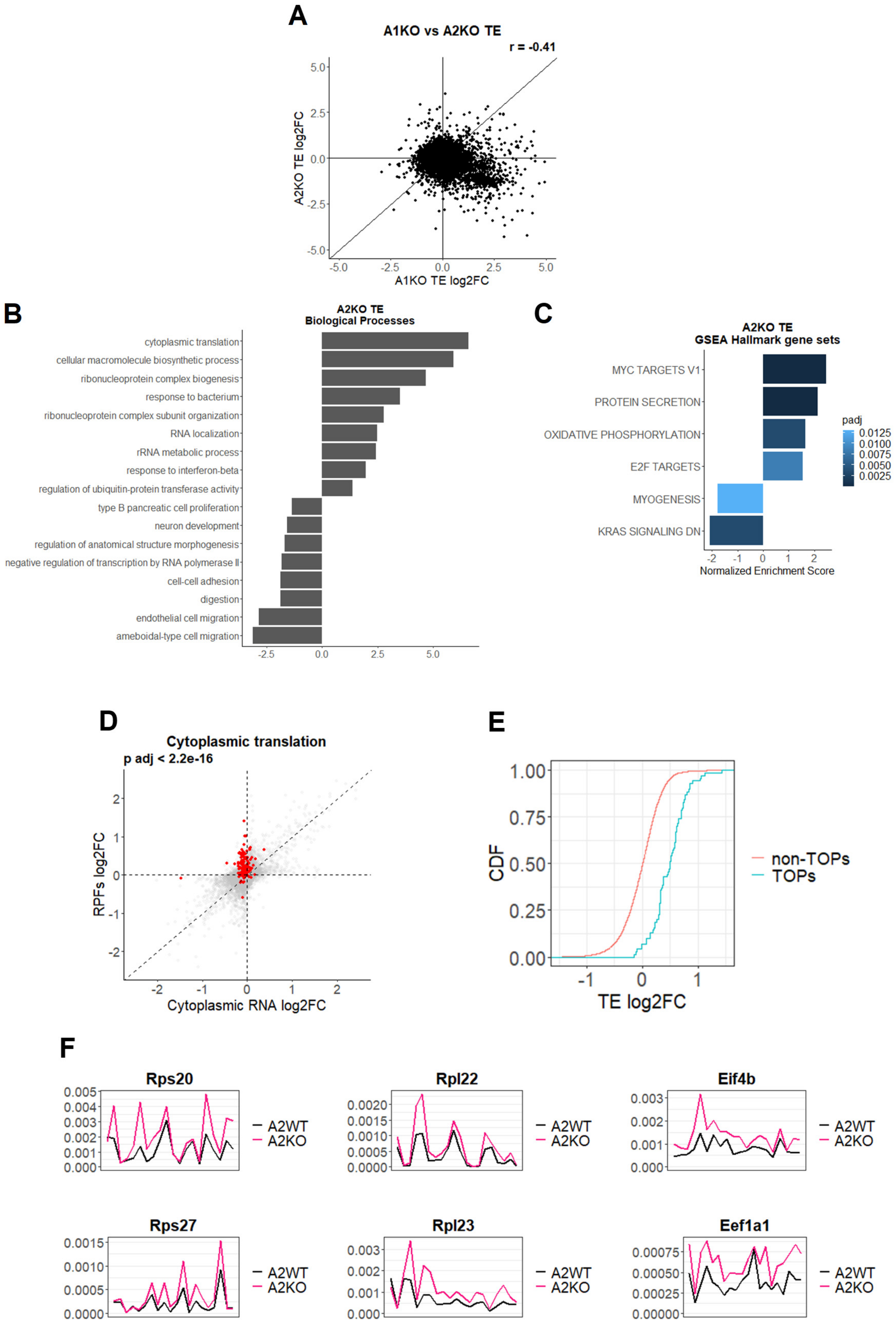


Fig. S8. (legend on following page)

**eIF4A2 represses translation of TOP mRNAs.** Ribosome profiling analysis following loss of eIF4A2 in *Apc*$^{fl/fl}$ *Kras*$^{G12D/+}$ small intestines, as in Fig. 4G-I and fig. S5. **(A)** Scatter plot comparing TE log2FCs following loss of eIF4A1 (A1KO) or eIF4A2 (A2KO), in *Apc*$^{fl/fl}$ *Kras*$^{G12D/+}$ small intestines. **(B)** GSEA using the biological processes as a reference and a ranked list of TE log2FCs as input. Redundant terms have been collapsed with rrvgo, using adjusted p-values as input for scores. Pathways enriched in genes translationally upregulated following eIF4A2 loss have positive values, and pathways enriched in downregulated genes have negative values. **(C)** GSEA using the Hallmark gene sets on a ranked list of TE log2FCs, with bar plot showing significantly enriched (padj < 0.05) Hallmark pathways in genes translationally altered following loss of eIF4A2. Pathways are ordered by normalized enrichment score (NES) and coloured by adjusted p-value. Pathways enriched in genes translationally upregulated following eIF4A2 loss have NES >0, and pathways enriched in downregulated genes have NES <0. **(D)** Overlaid TE scatter plots (from Fig. 4G), with genes from the "Cytoplasmic Translation" GSEA gene list coloured in red. Adjusted p-value was calculated with the fgsea R package. **(E)** Cumulative distribution function (CDF) plot showing the TE log2FC for previously described TOP mRNAs (*95*) and all remaining non-TOP mRNAs. **(F)** Meta plots depicting the normalised ribosome occupancy, across the length of the CDS, of the indicated TOP mRNAs.

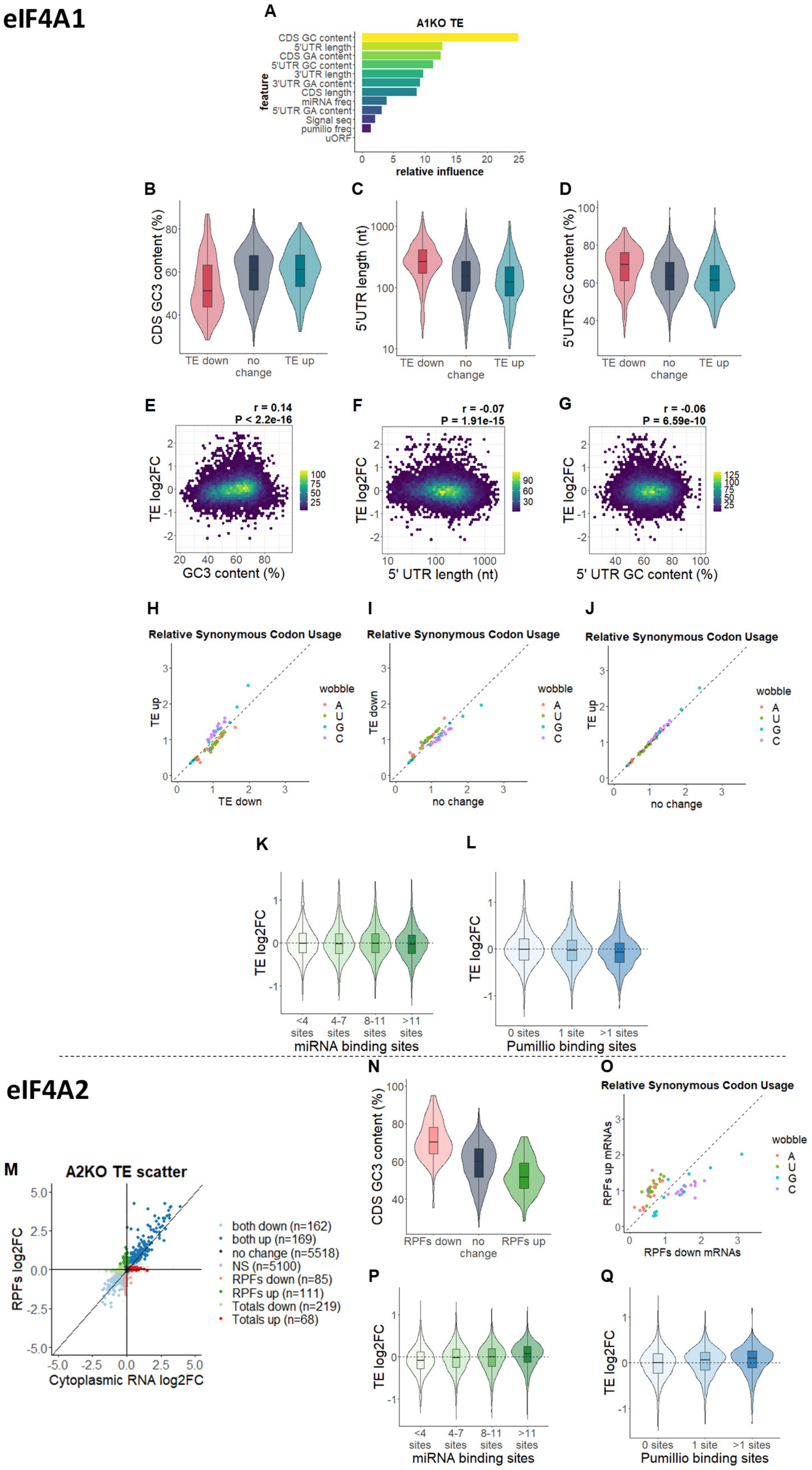


**Fig. S9. (legend on following page)**

**Feature properties of eIF4A1 and eIF4A2 dependent mRNAs. (A-L)** Analysis of the mRNA feature properties that best explain translational changes following loss of eIF4A1 in *Apc*$^{fl/fl}$ *Kras*$^{G12D/+}$ small intestines. (A) Relative influence scores for all feature properties, used within a gradient boosting approach, to predict which mRNA features contribute most towards TE log2FCs. (B to D) Violin with overlaid boxplots comparing GC3, 5'UTR length, and 5'UTR GC content between mRNAs within the A1KO TE "up/down" and "no change" groups from Fig. 4C. All comparisons have an adjusted p-value < 0.05, except "TE up" vs "no change" in GC3 content (p=0.48) and 5'UTR GC content (p=0.16). (E to G) Density scatter plots comparing GC3, 5'UTR length, and 5'UTR GC content against TE log2FCs. r- and p-values were obtained from a Pearson correlation test. Colour intensity scale indicates the number of transcripts within each hexagon, which is denoted in the legend. (H to J) Mean relative synonymous codon usage between mRNAs within the A1KO "TE down", "TE up" or "no change" groups, colour-coded by the wobble position of each codon. (K and L) Violin with overlaid boxplots showing TE log2FCs for mRNAs grouped by the number of predicted miRNA (K) or Pumilio (L) binding sites. All comparisons within (K) have an adjusted p-value > 0.05, except >11 sites vs either 4–7 sites (p=0.006) or 8–11 sites (p=0.005). All comparisons within (L) have an adjusted p-value < 0.05 except 0-sites vs 1-site (p=0.42). **(M-Q)** Analysis of the mRNA feature properties that best explain translational changes following loss of eIF4A2 in *Apc*$^{fl/fl}$ *Kras*$^{G12D/+}$ small intestines. (M) TE scatter plot depicting the RPF and total RNA log2FCs, coloured depending on whether the mRNA is differentially expressed in just the RPFs, or just the total cytoplasmic RNAs, or both. The "RPFs up" and "RPFs down" mRNAs are differentially up- or down-regulated in the RPFs (RPFs up, padj < 0.1 and log2FC > 0.2; RPFs down, padj < 0.1 and log2FC < -0.2), but are not differentially expressed in the same direction in the cytoplasmic RNA (RPFs up, padj ≥ 0.5 or log2FC < -0.2; RPFs down, padj ≥ 0.5 or log2FC > 0.2). The "Totals up" or "Totals down" mRNAs are as above but are differentially expressed in the cytoplasmic RNA but not in the same direction in the RPFs. The "both up" or "both down" mRNAs are differentially expressed in both the RPFs and the cytoplasmic RNA (padj < 0.1 and both up, log2FC > 0.2; both down, log2FC < -0.2). The "no change" mRNAs have padj ≥ 0.5 in both the RPFs and the cytoplasmic RNA and all remaining mRNAs are deemed non-significant (NS). (N) Violin with overlaid boxplot comparing GC3 content between mRNAs within the A2KO "RPFs up", "RPFs down" and "no change" groups from (M). All comparisons have an adjusted p-value < 0.05. (O) Mean relative synonymous codon usage between A2KO "RPFs up" and "RPFs down" mRNA, colour-coded by the wobble position of each codon. (P to Q) Violin with overlaid boxplots showing TE log2FCs for mRNAs grouped by the number of predicted miRNA (P) or Pumilio (Q) binding sites. All comparisons within (P) have adjusted p-values < 0.05, except between 4–7 and 8–11 sites (p=0.98). All comparisons within (Q) have adjusted p-values < 0.05, except between 1 site and >1 site (p=0.14). To test for statistical significance in B-D, K-L, N and P-Q, one-way ANOVA tests were performed with Tukey's multiple comparisons to compare all means.

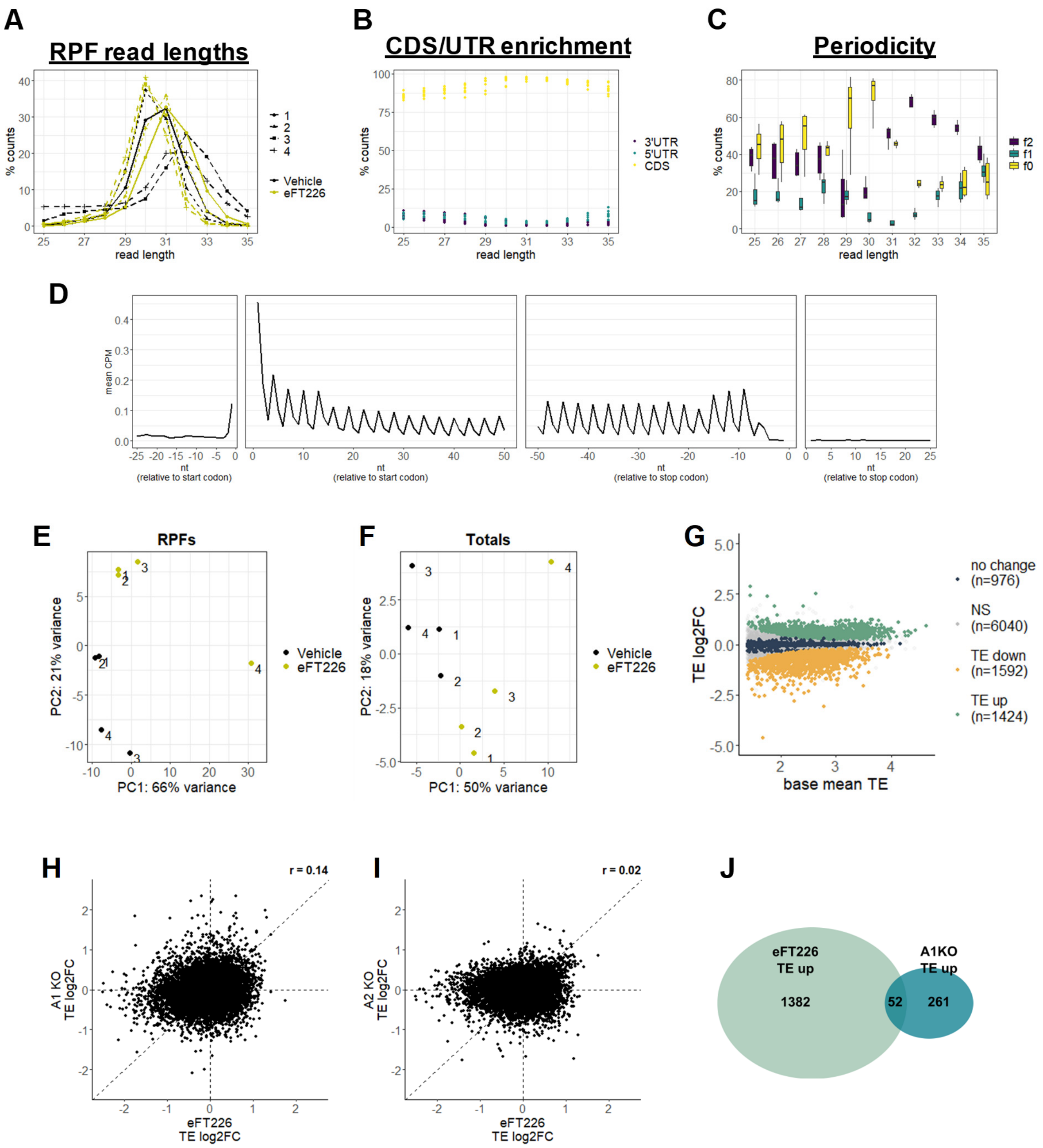


**Fig. S10.**

**Quality control of ribosome-profiling in small intestines following eFT226 treatment.** Vehicle/eFT226 = *VillinCreER*$^{T2}$ *Apc*$^{fl/fl}$ *Kras*$^{G12D/+}$ treated with vehicle/eFT226 2 h prior to sampling (n=4). **(A)** Ribosome protected fragments (RPF) read length distribution for indicated samples. **(B)** Percentage of protein-coding RPF reads that aligned to either the 5'UTR, CDS, or 3'UTR, separated by read length. **(C)** Boxplot depicting the percentage of RPF reads within each frame of the coding sequence, separated by read length. Read lengths 29–34 were taken forward for downstream analysis as these represented the highest CDS:3'UTR ratio of reads and best periodicity. **(D)** Average counts per million (CPM) at the indicated positions, up and down-stream of the start and stop codons, after applying an offset of 12 nt for read lengths 29–30 and

13 for read lengths 31–34. **(E-F)** Principal component analysis (PCA) plot of all RPF (E) and cytoplasmic RNA (F) samples. **(G)** Translational efficiency (TE) MA plot. TE log2FC = RPFs log2FC – Cytoplasmic RNA log2FC. Base mean TE is the mean TE (normalised RPFs - normalised cytoplasmic RNA) across all samples, as calculated with DESeq2. **(H** and **I)** Scatter plots comparing TE log2FCs following loss of either eIF4A1 (H) or eIF4A2 (I) versus eFT226 treatment. r-values were calculated with a Pearson correlation test. **(J)** Venn diagram depicting the overlap of mRNAs within the TE “up” groups, following eFT226 treatment or loss of eIF4A1.

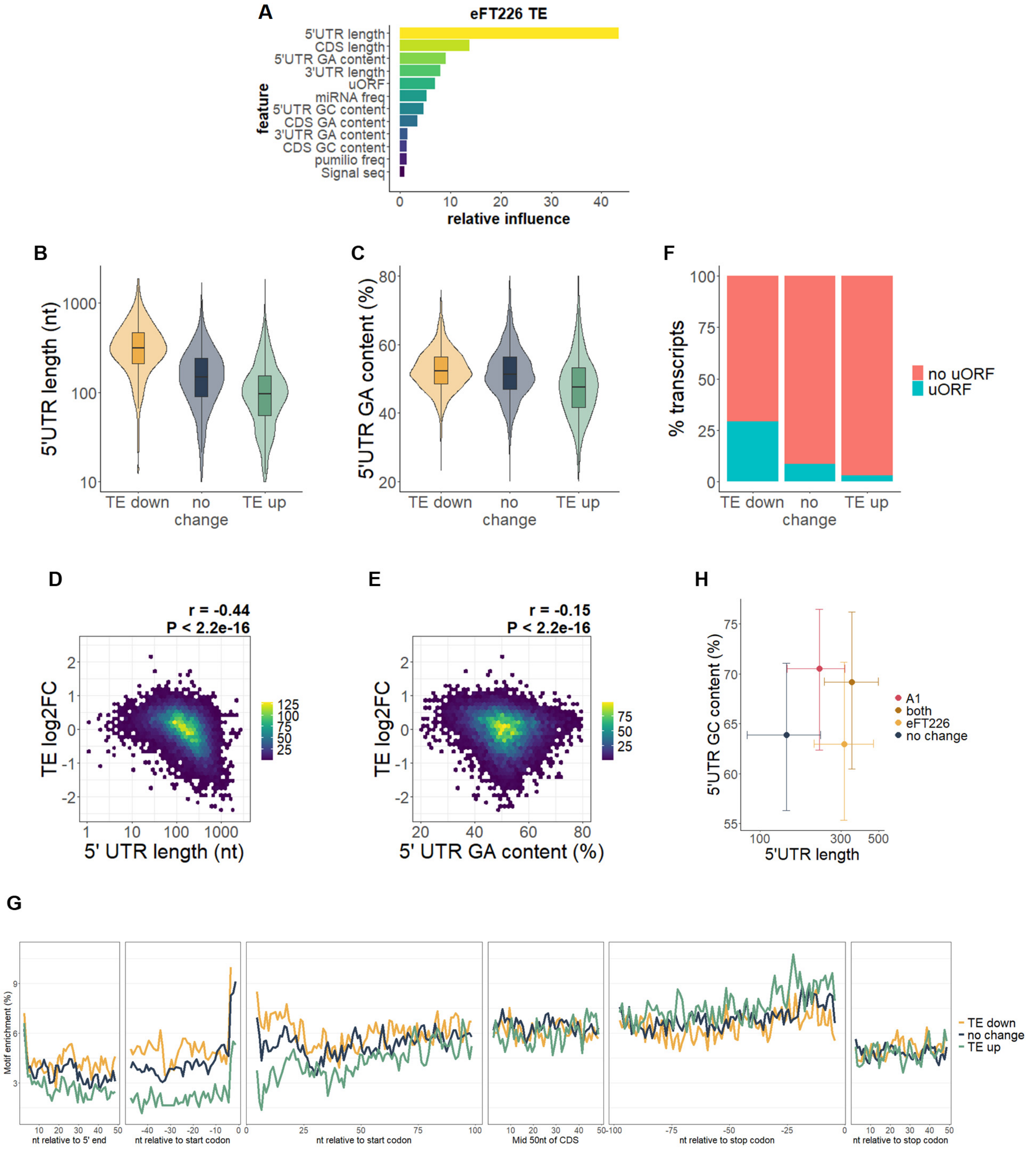


**Fig. S11.**

**eFT226 treatment clamps eIF4A onto purine-rich motifs.** Analysis of the mRNA feature properties that best explain translational changes following eFT226 treatment in *Apc*$^{fl/fl}$ *Kras*$^{G12D/+}$ small intestines. **(A)** Relative influence scores for all feature properties, used within a gradient boosting approach, to predict which mRNA features contribute most towards TE log2FCs. **(B and C)** Violin with overlaid boxplots comparing 5’UTR length and GA content between mRNAs within the “TE up”, “TE down” and “no change” groups from Fig. 5F. All comparisons have an adjusted p-value < 0.05. **(D and E)** Density scatter plots comparing 5’UTR length and GA content against TE log2FC following eFT226 treatment. r- and p-values were obtained from a Pearson correlation test. Colour intensity scale indicates the number of transcripts within each hexagon, which is denoted in the legend. **(F)** Bar chart depicting the percentage of transcripts with annotated uORFs (*92*) from the eFT226-treated TE groups. **(G)** Mean positional enrichment of 8 GA tetramers previously shown to have the highest affinity for eIF4A binding (*63*), across the length of all transcripts within the depicted groups. **(H)** Comparison of 5'UTR length and GC content of the mRNAs from Fig. 5H. Dots, median; bars, interquartile range. All 5'UTR length comparisons have an adjusted p-value < 0.05 except eFT226 vs both (p=0.49). All 5'UTR GC content comparisons have an adjusted p-value < 0.05 except A1 vs both (p=0.65) and "no change" vs eFT226 (p=0.46). To test for statistical significance, one-way ANOVA was performed with Tukey's multiple comparisons to compare all means.

The data in this figure show that 5'UTR length has the strongest influence on TE log2FC, following eFT226 treatment, with a stronger correlation than following loss of eIF4A1 (compare panel (C) with fig. S9F). Importantly, the 5'UTR GA content and the presence of uORFs have a stronger influence than 5'UTR GC content. Furthermore, the enrichment of GA motifs is strongest towards the end of the 5'UTR and within the first 25 nt of the CDS. This suggests that eFT226 represses translation by clamping eIF4A onto purine-rich motifs, as opposed to altering 5'UTR structure, as is assumed to occur following loss of eIF4A1.

By comparing 5'UTR length and GC content, within the groups of mRNAs from Fig. 5H, we find that eIF4A1-dependent mRNAs have longer 5'UTRs, but eFT226-sensitive mRNAs are longer still. However, eIF4A1-dependent mRNAs have more GC-rich 5'UTRs, irrespective of whether they are also sensitive to eFT226 treatment. This suggests that there are two separate mechanisms for repressing translation, following loss of eIF4A1 or eFT226 treatment, with both processes reliant on 5'UTR length. eIF4A1-dependency seems to be driven by both 5'UTR GC content and length, which has been shown previously to increase the likelihood of secondary structures that are sensitive to eIF4A1 activity (*14*). In contrast, eFT226 treatment causes eIF4A to clamp onto mRNAs within purine-rich motifs, increasing the utilisation of inhibitory uORFs. In addition, the mRNAs most sensitive to eFT226 treatment contain longer 5'UTRs, which could increase the likelihood of the occurrence of a purine-rich motif downstream of a non-canonical start site.

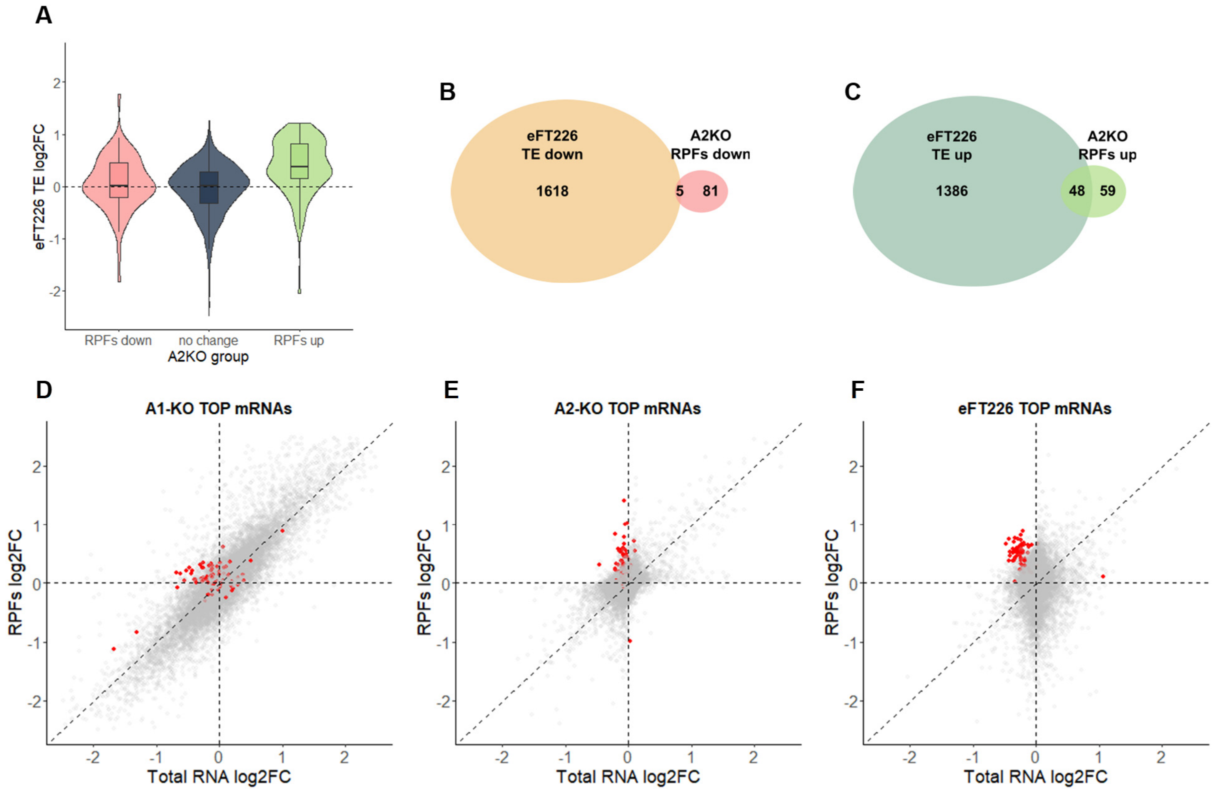


**Fig. S12.**

**Overlap of translational targets of eFT226 treatment and loss of eIF4A2. (A)** Violin with overlaid boxplots depicting the TE log2FC following eFT226 treatment, for the A2KO RPFs only groups from fig. S9M. To test for statistical significance, an ANOVA was performed with Tukey's multiple comparisons to compare all means. All comparisons had an adjusted p-value < 0.05, except for "RPFs down" versus "no change" (p=0.3). **(B** and **C)** Venn diagrams depicting the overlap in mRNAs between the eFT226 TE groups from Fig 4F and the A2KO RPFs only groups. This shows that 44.9% of the mRNAs which go up translationally following loss of eIF4A2, also increase in translational efficiency with eFT226 treatment, while the same is only true for 5.8% of the mRNAs that decrease in translational efficiency. It should be noted that most of the mRNAs that increase translationally in both conditions (29/48) are TOPs. **(D** to **F)** TE scatter plots for eIF4A1 KO (D), eIF4A2 KO (E), and eFT226 treatment (F), overlaid with previously described TOPs (*95*). The TE of all TOP mRNAs increases following loss of eIF4A2, at the RPF level, with little change at the total RNA level. On the other hand, the TE of TOP mRNAs also increases following loss of eIF4A1, but this is driven through a decrease in total RNA level, with no concomitant change in the RPF level. Interestingly, following eFT226 treatment, TOP mRNAs are increased at the RPF level and are downregulated at the total RNA level, which is consistent with this drug inhibiting both eIF4A paralogues, resulting in an additive effect on TOP mRNAs.

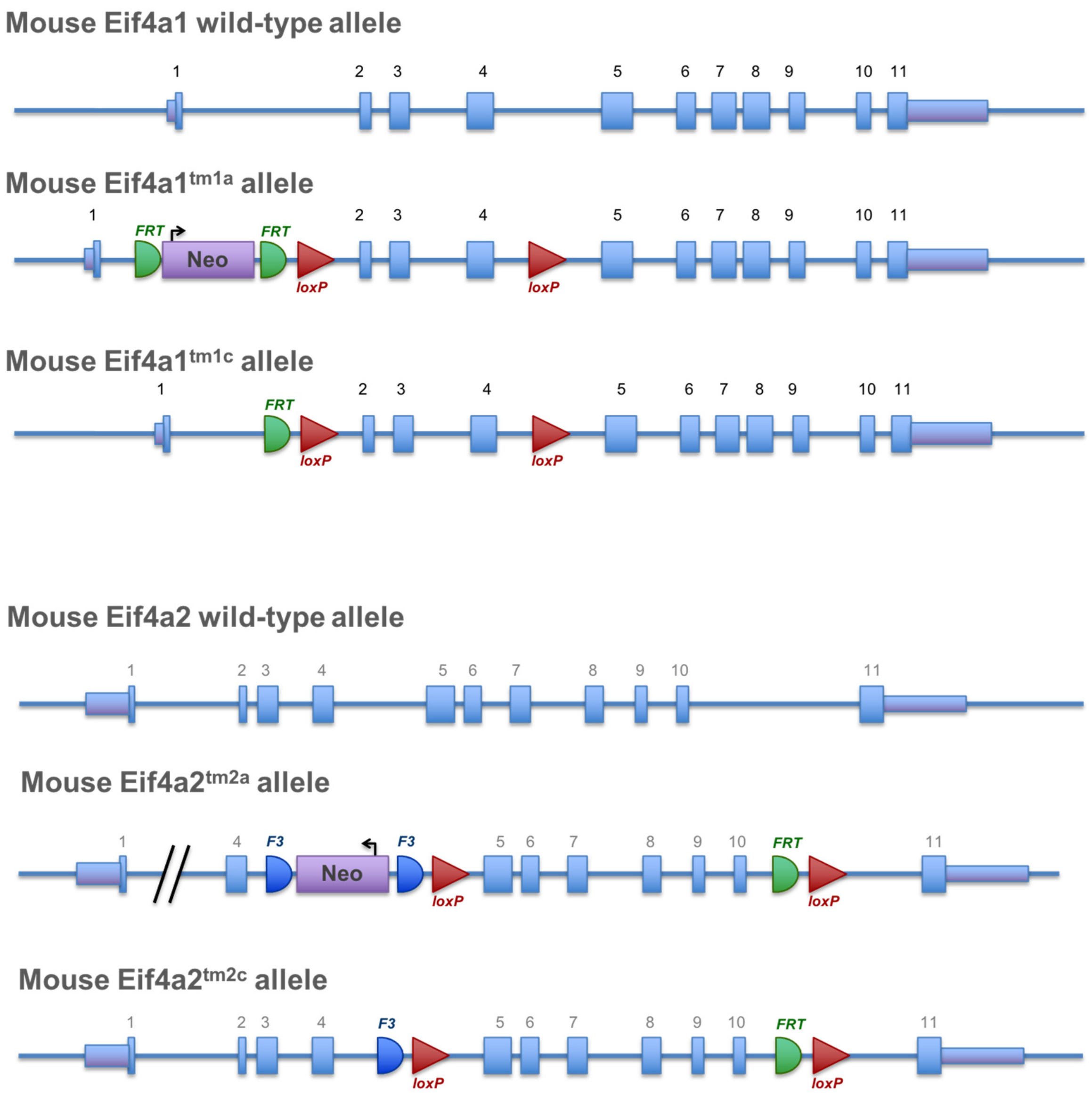


**Fig. S13.**
**Diagrammatic representation of the *Eif4a1^fl^* and *Eif4a2^fl^* alleles used for conditional knockout. (A)** Generation of the *Eif4a1^fl^* allele (see methods for further details). **(B)** Generation of the *Eif4a2^fl^* allele (see methods for further details).

**Data S1. (separate file)**

Output from fgsea using log2 fold changes in translational efficiency (TE log2FC) following loss of either eIF4A1 (A1KO) or eIF4A2 (A2KO) in *Apc*$^{fl/fl}$ *Kras*$^{G12D/+}$ mouse small intestines, as input and either the GSEA mouse specific Curated, Hallmark or Biological Processes terms as references.
